# The mechanisms of catalysis and ligand binding for the SARS-CoV-2 NSP3 macrodomain from neutron and X-ray diffraction at room temperature

**DOI:** 10.1101/2022.02.07.479477

**Authors:** Galen J. Correy, Daniel W. Kneller, Gwyndalyn Phillips, Swati Pant, Silvia Russi, Aina E. Cohen, George Meigs, James M. Holton, Stefan Gahbauer, Michael C. Thompson, Alan Ashworth, Leighton Coates, Andrey Kovalevsky, Flora Meilleur, James S. Fraser

## Abstract

The NSP3 macrodomain of SARS CoV 2 (Mac1) removes ADP-ribosylation post-translational modifications, playing a key role in the immune evasion capabilities of the virus responsible for the COVID-19 pandemic. Here, we determined neutron and X-ray crystal structures of the SARS-CoV-2 NSP3 macrodomain using multiple crystal forms, temperatures, and pHs, across the apo and ADP-ribose-bound states. We characterize extensive solvation in the Mac1 active site, and visualize how water networks reorganize upon binding of ADP-ribose and non-native ligands, inspiring strategies for displacing waters to increase potency of Mac1 inhibitors. Determining the precise orientations of active site water molecules and the protonation states of key catalytic site residues by neutron crystallography suggests a catalytic mechanism for coronavirus macrodomains distinct from the substrate-assisted mechanism proposed for human MacroD2. These data provoke a re-evaluation of macrodomain catalytic mechanisms and will guide the optimization of Mac1 inhibitors.

## Introduction

Viral infection triggers the release of interferons, a group of secreted cytokines that are key components of the innate immune response (*1*–*4*). Among the several hundred interferon-stimulated genes are poly-(ADP-ribose) polymerases (PARPs), a family of enzymes that catalyze the transfer of ADP-ribose from NAD+ to proteins as a post-translational modification. Both mono- and poly-ADP-ribosylation are important for signal transduction of the interferon response (*5*–*7*). In SARS-CoV-2 infection, ADP-ribosylation is mediated by PARP9 and its binding partner DTX3L (*8*). To counteract this host defense mechanism, SARS-CoV-2 encodes a macrodomain, Mac1, with mono-(ADP-ribosyl)-hydrolase activity that reverses the protective effect of ADP-ribosylation (*9*) (**Fig. 1**A). Importantly, an active site mutation that inactivates Mac1 has been shown to render SARS-CoV non-lethal in a mouse model of viral infection (*10*) and interferon stimulated cells can clear SARS-CoV virus bearing that active site mutation, but not WT Mac1. Accordingly, Mac1 is a potential therapeutic target for small molecule inhibitors. Importantly, the Mac1 active site is highly conserved between SARS-CoV, SARS-CoV-2 and MERS, which raises the possibility that a Mac1 inhibitor could be used as a broad spectrum antiviral against coronaviruses (*11*).

**Fig. 1.**
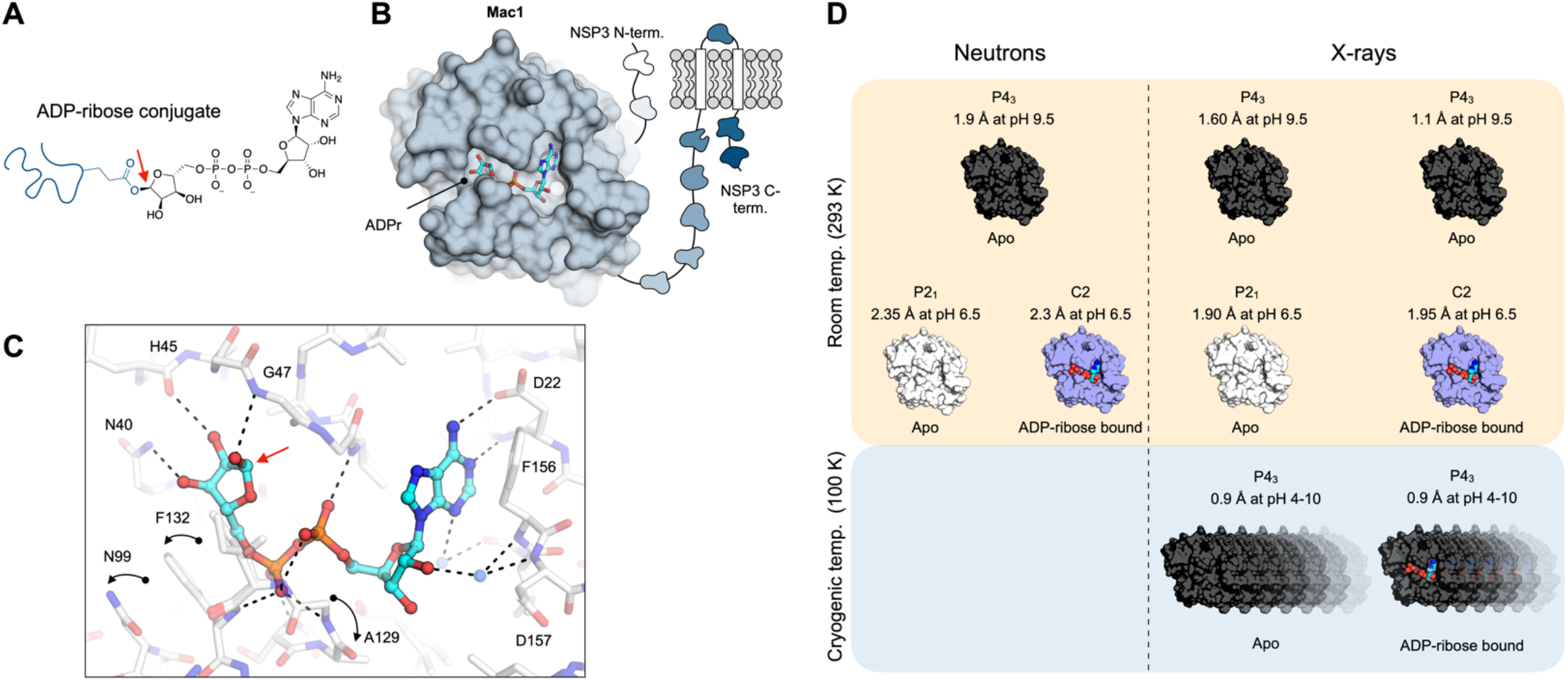
The NSP3 macrodomain (Mac1) reverses mono-ADP-ribosylation. (**A**) Chemical structure of ADPr showing the C1’’ covalent attachment point with a red arrow. (**B**) Cartoon of the multi-domain NSP3 showing Mac1 with ADPr bound in the active site (PDB code 7KQP). (**C**) Structure of ADPr bound in the macrodomain active site (PDB code 7KQP) with the changes in protein structure upon ADPr binding indicated with black arrows. (**D**) Summary of the crystal structures reported in this work.

The SARS-CoV-2 macrodomain is encoded as a domain of non-structural protein 3 (NSP3), a large multi-domain protein found in all coronaviruses (*12*) (**Fig. 1**B). Mac1 adopts an α/β/α-sandwich fold and binds ADPr through an extensive network of hydrogen bonds in a well-defined cleft (**Fig. 1**B/C) (*9, 13*). Several conformational changes are stabilized upon ADPr binding, including rotation of Phe132 to accommodate the terminal ribose of ADPr, and two peptide flips to bind the diphosphate portion of ADPr (*14*). Although the structural determinants of ADPr-binding are well established, the catalytic mechanism of Mac1 is unknown. Based on structural homology to the human macrodomain hMacroD2 (*15*), a substrate-assisted mechanism has been proposed for viral macrodomains (*16*), however, this has not been confirmed experimentally. Determining the precise orientations of active site water molecules and the protonation states of key catalytic site residues can discriminate whether the proposed water nucleophile is appropriately orientated for the hydrolysis reaction in the ADPr-bound state or if other catalytic mechanisms should be considered.

Although X-ray crystallography can provide the high resolution structural information, hydrogen atoms are not normally observed in X-ray diffraction experiments (*17*). This means that key information may be obscured, including residue protonation states, hydrogen bond networks and water orientations. Neutron crystallography allows hydrogens to be visualized at modest resolutions (<2.5 Å) (*18*–*21*), which can reveal details missing from X-ray structures. In practice, hydrogen is typically exchanged for deuterium in protein samples for neutron crystallography, because protons have a small negative neutron scattering length, as well as a large incoherent cross section, whereas deuterium has a comparatively large positive neutron scattering length and small incoherent cross section (*22*). The low brightness of neutron sources means that the volume of crystals required is 100-1000-fold larger than crystals that are suitable for experiments at synchrotron light sources (*23*). Because protein crystals rarely grow to this size, relatively few drug targets have been structurally characterized by neutron crystallography (*24*–*26*).

In addition to informing catalytic mechanisms, neutron crystallography can augment structure-based drug design, which depends upon an accurate model of the shape, electrostatic potential, solvation and flexibility of the target site (*27*–*29*). We recently screened more than 2500 fragments against Mac1 using X-ray crystallography (*14*). Structures of 234 fragments bound to Mac1 were determined, including almost 200 in the active site. This screen has given the community an extensive set of chemically diverse starting points for fragment-based ligand discovery against Mac1. As expected based on their low molecular weight, the fragments bound Mac1 weakly (*14*). Efforts have been assisted by the development of Mac1-specific biophysical and biochemical assays (*8, 30, 31*). The task of converting fragments into potent and selective Mac1 inhibitors will be helped by a more detailed understanding of the physical chemical features of the Mac1 active site.

Here, we report a series of Mac1 crystal structures determined using neutron and X-ray diffraction (**Fig. 1**D). Two neutron structures of the apo enzyme were determined from distinct crystal forms grown at pH 6.5 and 9.5, and a neutron structure of the ADPr-Mac1 complex was determined at pH 6.5. Protonation states and active site hydrogen bond networks were characterized in the three structures, revealing the molecular basis for functionally relevant flexibility in active site loops and proton locations relevant to the catalytic mechanism. We mapped water networks in the active site, including water orientations from neutron diffraction, and showed that the networks were robust to changes in crystal packing, temperatures and pH. The water networks reorganize upon ADPr binding, with a network of tightly bound water molecules acting as protein-ligand bridges. Finally, we show how these same tightly bound water molecules are co-opted by fragment binding, with implications for inhibitor design. These results advance our knowledge about the structure and function of viral macrodomains, and provide a structural resource to guide the design of macrodomain inhibitors as SARS-CoV-2 antiviral therapeutics.

## Results

### Room temperature Mac1 structures from distinct crystal forms at different pHs

#### Mac1 neutron structure at pH 9.5 (1.9 Å, P4_3_ crystals)

We grew neutron diffraction quality crystals at pH 9.5 using a Mac1 construct that crystallizes in the P4_3_ space group (*14*) (**Fig. 2**A). Hydrogen/deuterium exchange (HDX) was achieved by growing crystals using solutions prepared with D_2_O, followed by soaking the crystal in a mother liquor prepared with D_2_O. Neutron diffraction data to 1.9 Å were collected at room temperature using the Macromolecular Neutron Diffractometer (MaNDi) at the Oak Ridge National Laboratory (*32*–*34*) (**Table S1**). To enable joint neutron/X-ray refinement in Phenix (*35*), we collected a 1.6 Å X-ray diffraction dataset from the same crystal using a rotating anode X-ray source (**Table S1**). To account for differences in HDX at different protein sites, both hydrogen and deuterium were modeled at exchangeable positions, and deuterium occupancies were automatically refined (*35*). After several cycles of refinement and model building, the *R*_work/_*R*_free_ values were 22.6/27.5% for the neutron data and 12.4/15.7% for X-ray data. In total, 203 D_2_O molecules were modeled across the two molecules in the asymmetric unit (ASU) (**Fig. 2**B). There was weak electron density for *N*-cyclohexyl-2-aminoethanesulfonic acid (CHES) in the adenosine site, with the CHES sulfonate interacting with the backbone nitrogens of Phe156/Asp157 (the oxyanion subsite), mimicking the bridging interactions between a water molecule and the proximal ribose of ADPr (**Fig. S1**). The atomic coordinates and structure factor amplitudes have been deposited in the Protein Data Bank with the accession code 7TX3.

**Fig. 2.**
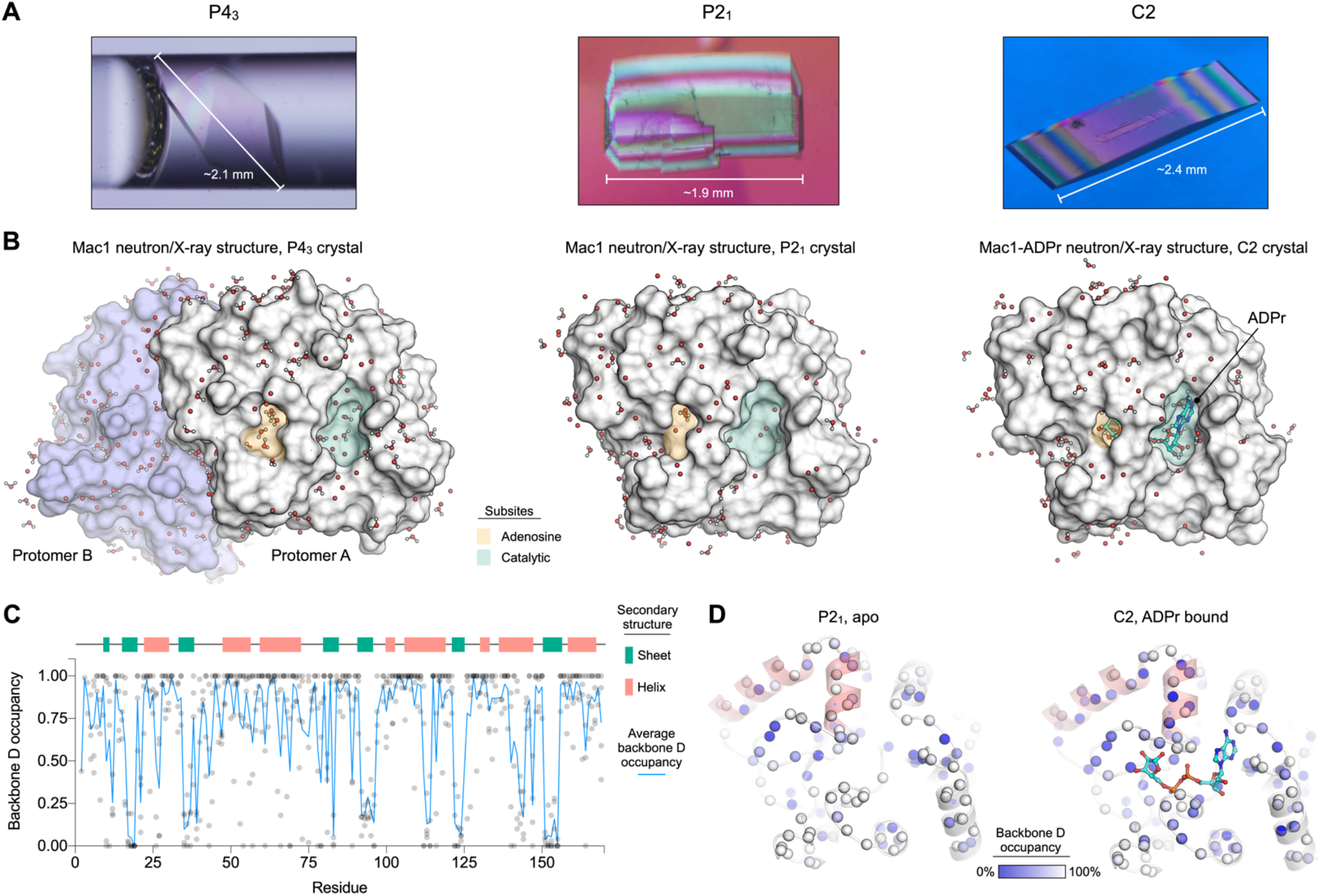
Crystal structures of Mac1 determined using neutron diffraction. (**A**) Neutron quality Mac1 crystals grown in the P4_3_, P2_1_ and C2 space groups. The P4_3_ crystal is shown in the quartz capillary used for data collection, while the P2_1_ and C2 crystals are shown prior to mounting. (**B**) Molecular surface showing Mac1 neutron structures. D_2_O molecules within 5 Å of the protein surface are shown with sticks/spheres. The adenosine and catalytic sites are shaded green and yellow respectively. (**C**) Plot showing the occupancy of backbone amide deuterium atoms in the three Mac1 crystal structures. (**D**) Backbone amide deuterium occupancy mapped onto the P2_1_ and C2 Mac1 structures. Backbone nitrogens are shown with blue spheres, colored by backbone D occupancy (blue=0%, white=100%). The helices composed of residues 50-70 are shaded red. The average backbone D occupancy for these helices was 82% in the P2_1_ structure and 52% in the C2 structure.

#### Mac1 X-ray structure at pH 9.5 (1.1 Å, P4_3_ crystals)

To obtain more structural information about Mac1 at room temperature, we collected higher resolution diffraction data using the P4_3_ crystals and synchrotron radiation (**Table S1**). The sensitivity of the P4_3_ crystals to radiation damage was assessed by collecting four datasets from similarly sized crystals (∼0.1 mm^3^), with the absorbed X-ray dose varied from 73 to 539 kGy. There was no evidence for general or specific radiation damage, as measured by per-image resolution, *R*_CP_, unit-cell parameters and side-chain B-factors (**Fig. S2**). The X-ray dose and dataset resolution were correlated, with the highest dose dataset having the highest resolution (1 Å, truncated to 1.1 Å to achieve ∼100% completeness). Consistent with radiation damage-free structures, the refined coordinates and B-factors from the four datasets were close to identical (Cα root-mean-square fluctuation (RMSF) < 0.1 Å, Pearson correlation coefficient (r) for Cα B-factors >0.98) (**Fig. S2**). The atomic coordinates and structure factor amplitudes from the four datasets have been deposited in the PDB with the accession codes 7TWF, 7TWG, 7TWH, 7TWI. Based on having similarly high resolution, but slightly lower Cα and side chain B-factors, the structure refined from the 290 kGy dataset was used for subsequent structural analysis (*R*_work_/*R*_free_ = 9.94/11.5%).

Alignment of the 1.1 Å X-ray structure with the 1.9/1.6 Å neutron/X-ray structure shows that the protein coordinates are nearly identical (Cα root-mean-square deviation (RMSD) <0.05 Å, Pearson r >0.93 for Cα B-factors) (**Fig. S3**). Although the crystals used for X-ray diffraction were grown in the same conditions as the neutron crystal (100 mM CHES pH 9.5, 34% PEG 3000), there was no evidence for CHES in the active site. This may be due to slight differences in how the crystals were handled. There were slightly more water molecules modeled in the 1.1 Å structure compared to the jointly refined neutron/X-ray structure (360 versus 317). Comparison of the waters between the two structures shows that their positions are conserved (**Fig. S3**). Of the 317 waters modeled in the 1.9/1.6 Å neutron/X-ray structure, 257 had a matching water within 0.5 Å in the 1.1 Å X-ray structure. The waters that were not conserved tended to have higher B-factors, indicating increased disorder (**Fig. S3**).

#### Mac1 neutron structure at pH 6.5 (2.3 Å, P2_1_ crystals)

To investigate structural differences in Mac1 across different crystal forms and different pHs, we also grew neutron-quality crystals at pH 6.5 using a construct that crystallized in the P2_1_ space group (*36*) (**Fig. 2**A/B). The difference between the two constructs was slight: the P2_1_ construct contained a Gly-Glu sequence immediately following the TEV cleavage site, while the P4_3_ construct had Ser-Met at the equivalent position. Hydrogen/deuterium exchange was performed in a similar manner to the P4_3_ crystals, and diffraction data was collected to 2.35 Å at MaNDi (**Table S1**). As with the P4_3_ crystals, an X-ray dataset was collected with the same crystal (**Table S1**), and joint X-ray/neutron refinement was performed with phenix.refine. The final *R*_work/_*R*_free_ values were 17.7/26.1% for the neutron data and 16.6/22.5% for X-ray data. Overall, the structure was similar to both monomers of the P4_3_ structure (Cα RMSD <0.2 Å). Although MES was bound at full occupancy in the previously reported X-ray structure using the same construct and similar crystallization conditions (*36*), there was no evidence for MES binding in the new structure. The atomic coordinates and structure factor amplitudes have been deposited in the PDB with the accession code 7TX4.

#### ADPr-bound Mac1 neutron structure at pH 6.5 (2.3 Å, C2 crystal)

To better understand the molecular basis for ADPr recognition by Mac1, we co-crystallized ADPr with the P2_1_ construct (**Fig. 2**A/B). Unlike the P4_3_ construct, where crystal packing prevents co-crystallization with ADPr (*14*), co-crystals with the P2_1_ construct grew readily. Neutron diffraction data was collected to 2.3 Å using the IMAGINE beamline at the High Flux Isotope Reactor (HFIR) at ORNL (*37*) and an X-ray dataset was collected to 1.95 Å using the same crystal. The space group of the co-crystal was C2, indicating that binding of ADPr had perturbed crystal packing. We performed joint neutron/X-ray refinement of the ADPr-Mac1 data using phenix.refine, with *R*_work/_/*R*_free_ values of 18.7/26.2% for the neutron data and 13.3/17.4% for X-ray data. There was clear positive density for ADPr bound in the Mac1 active site in the mF_O_-DF_C_ electron density map (**Fig. S1**), and the ADPr occupancy was refined to 93%. The terminal ribose of ADPr is bound in the β-configuration, with the C1’’ hydroxyl hydrogen bonded to the backbone nitrogen of Gly48. Alignment of the Mac1-ADPr neutron/X-ray structure with the three apo structures shows that the structures are highly similar (Cα RMSD <0.3 Å) (**Fig. S3**). Structural differences include the peptide flips of Gly48 and Ala129 that allow two new hydrogen bonds with the diphosphate portion of ADPr, and a coupled conformational change in the Phe132 and Asn99 side-chains to accommodate the terminal ribose of ADPr (**Fig. S1**). These changes have been documented across several crystal forms (*9, 13, 36, 38*). The main difference with the ADPr-bound structure obtained with the P4_3_ crystal form is that crystal packing interactions prevent the flip of Gly48, forcing ADPr to adopt the α configuration (*14*). The atomic coordinates and structure factor amplitudes for the ADPr-bound neutron/X-ray structure have been deposited in the PDB with the accession code 7TX5.

#### Features of deuteration between the different crystal structures

Next, we examined whether the refined deuterium occupancy in the neutron structures could reveal information about Mac1 structure and dynamics (*39*–*41*). Although deuterium occupancy was refined at all exchangeable positions, we restricted our analysis to backbone amide deteriums. For the P4_3_ crystal, the average backbone nitrogen deuterium occupancy was 78 and 76% for the two monomers in the ASU, and there was good agreement between the refined deuterium occupancies (Pearson r = 0.87) (**Fig. S4**). Because the monomers have unique lattice contacts, this suggests that the differences in HDX are caused by structural features of Mac1, not merely differences in crystal packing. Although the average backbone nitrogen deuterium occupancy in the P2_1_ and C2 crystals were slightly lower (at 69% and 57%, respectively), there was still good agreement between the refined backbone amide deuterium occupancy (**Fig. S4**). In all four monomers, backbone deuteriums with low occupancy tended to be located on the buried β-strands (**Fig. 2**C, **Fig. S4**), which is consistent with the protected nature of backbone amides in β-sheets (*39*). The protective effect was pronounced in the solvent exposed β-strand that includes residues 80-84, where the refined D occupancies alternate between high and low, depending on whether the backbone amide is solvent exposed or not (**Fig. S4**). Compared to the P4_3_ crystals, there were more positions that were refined with 0% occupancy in the ADPr-bound structure determined using C2 crystals (**Fig. S4**). This included a cluster of residues on the two α-helices composed of residues 50-70 (**Fig. 2**D). This may reflect increased protection from HDX caused by stabilization of the Gly46-48 loop in the ADPr-bound structure. However, comparing deuteration between crystal forms is complicated by differences in crystal packing, different pH of crystallization and differences in detuteration procedure. For example, the α-helix composed of residues 138-145 has relatively low HDX in the P2_1_ and C2 crystals, which may be connected to substantial differences in local crystal packing (**Fig. S4**). Despite the possible connection between lattice contacts and HDX, the refined deuterium occupancies were only weakly correlated with crystallographic B-factors (**Fig. S4**).

### Assignment of Mac1 protonation states at pH 6.5 and 9.5

#### Histidines

Histidine can adopt multiple protonation states and tautomers at physiological pH, which allows it to play diverse roles in protein structure and function. Mac1 contains four histidines that are exposed to the bulk solvent, and two that are buried (**Fig. 3**A). We assigned histidine protonation states based on F_O_-F_C_ neutron scattering length (NSL) difference density maps (**Fig. 3**B/C, **Fig. S5**). The histidine protonation states were the same across both crystal forms, except for histidines 86, 119 and 138, which were singly protonated in the P4_3_ structure, but doubly protonated in the P2_1_ structure (**Fig. 3**C). The importance of experimentally determining protonation states is highlighted by the mismatch between the protonation states predicted by the Reduce program (*42*) and the states assigned using neutron crystallography (**Fig. 3**C). Although the protonation states of the two buried histidines, His45 and His94, were assigned correctly by Reduce, there was a discrepancy in the protonation states for the remaining histidines in at least one of the P4_3_ protomers (**Fig. 3**C). To test whether histidine protonation states could be assigned from the previously published high resolution X-ray structures of Mac1 (*14*), we calculated F_O_-F_C_ electron density difference maps from the 0.86 Å X-ray structure determined using the P4_3_ crystals (PDB code 7KQO), and the 0.77 Å structure determined using C2 crystals (PDB code 7KR0). Although the protonation state of His45 and His94 could be assigned based on the electron density maps, the protonation states of the other residues were ambiguous (**Fig. S5**). This highlights the power of neutron crystallography for assigning histidine protonation states, even when X-ray data is available at ultra-high resolution.

**Fig. 3.**
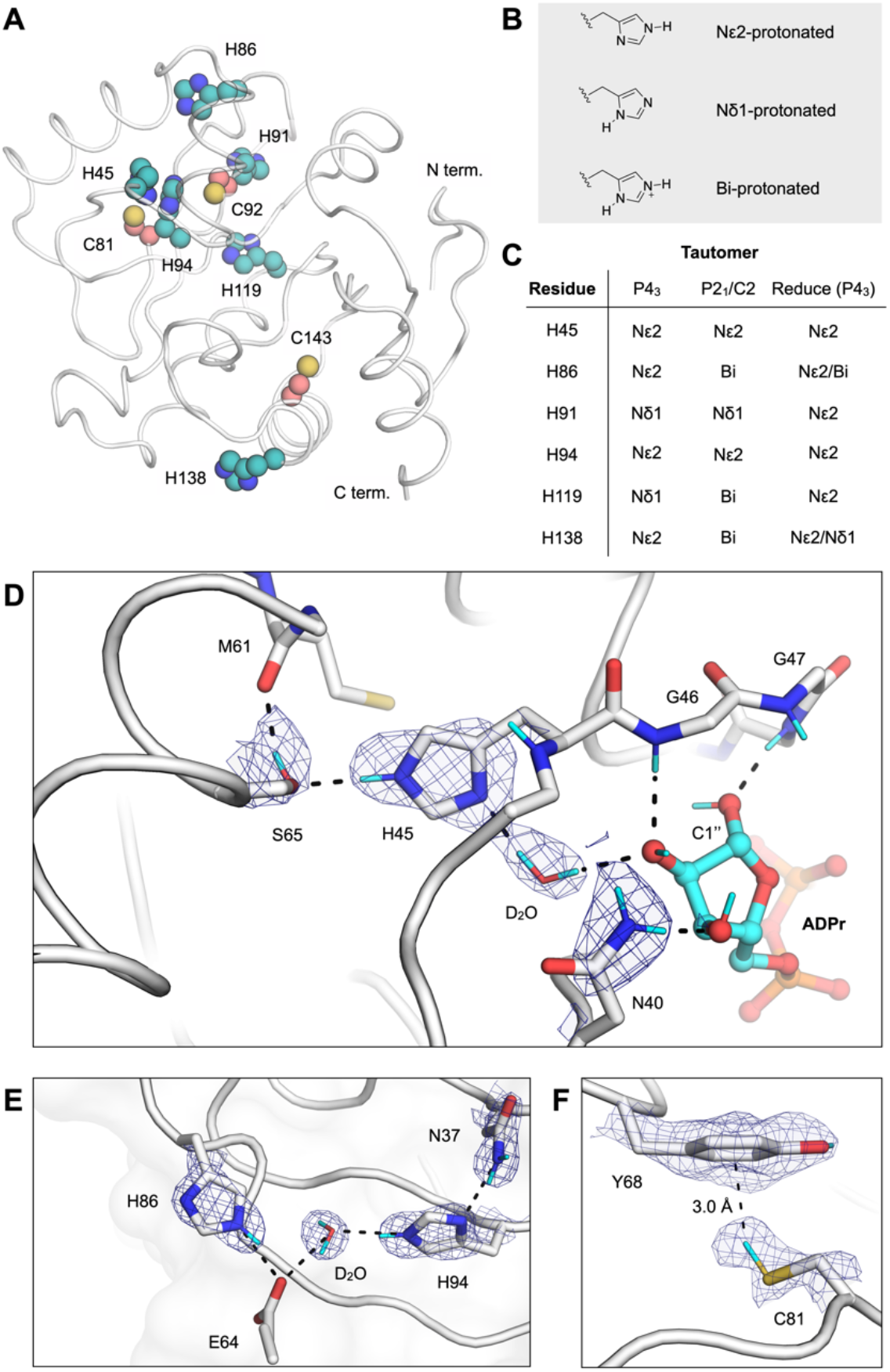
Protonation states of Mac1 histidine residues assigned by neutron diffraction. (**A**) The location of histidine (teal spheres) and cysteine (salmon spheres) residues mapped onto the Mac1 structure (PDB code 7KQO). (**B**) Chemical structures showing the three possible protonation states of histidine. (**C**) Histidine protonation states assigned based on NSL density maps (maps are shown in **Fig. S5**). The tautomers assigned to the high resolution P4_3_ X-ray structure (PDB code 7KQO) by the program Reduce are also shown. (**D**) NSL density maps reveal the hydrogen bond network connecting His45 and ADPr in the C2 structure (PDB code 7TX5). The protein is shown with a white cartoon/stick representation and the 2mF_O_-DF_C_ NSL density map is shown with blue mesh (contoured at 2.5 σ). (**E**) An extensive hydrogen bond network connects the Asn37 side chain with a surface histidine (His86). The P4_3_ structure (protomer A, PDB code 7TX3) and the corresponding 2mF_O_-DF_C_ NSL density map is shown with blue mesh (contoured at 2.5 σ). (**F**) An aromatic-thiol bond was observed between Tyr68 and Cys81 in protomer A of the P4_3_ structure (PDB code 7TX3). The 2mF_O_-DF_C_ NSL density map is shown with blue mesh (contoured at 1 σ).

Next, we examined the hydrogen bonding networks around the buried histidines. His45 is singly protonated on the Nε2 nitrogen, and hydrogen bonds to the Ser65 side chain and the backbone carbonyl of Gln62 (**Fig. 3**D). The neutron structure of Mac1 with ADPr bound shows that this network extends to the active site via a D_2_O molecule, which forms a hydrogen bond with the C2’’ hydroxyl of the terminal ribose (**Fig. 3**D). It has previously been shown that mutation of His45 to an alanine substantially decreases Mac1 activity (*16*). This water mediated network provides a potential link between His45 and Mac1 activity: His45 could participate either through stabilization of the terminal ribose in a catalytically productive conformation, as proposed previously(*15*), or through the action of His45 as a general base in the hydrolysis reaction. The other buried histidine, His94, participates in an extensive hydrogen bond network connecting the core of the Mac1 to its surface (**Fig. 3**E). Asn37 is hydrogen bonded to the singly protonated His94, and the network extends to His86 via a D_2_O molecule and the Glu64 backbone (**Fig. 3**E). The residues comprising these two networks are conserved in the SARS-CoV and MERS macrodomains (*16*).

#### Cysteines

The p*K*_a_ of the cysteine thiol (8.5) means that it often acts as a nucleophile in enzyme-catalyzed hydrolysis reactions. Mac1 has three buried cysteines, however, none are located in the active site, and they are unlikely to play a direct role in ADPr hydrolysis (**Fig. 3**A). To determine the protonation states of the Mac1 cysteines in the P4_3_ and P2_1_ crystal structures, we modeled the protonated and deprotonated forms and calculated F_O_-F_C_ NSL density difference maps (**Fig. S5**). Assigning the cysteine protonation states was difficult. There was a large positive peak placing the Cys81 thiol deuterium atom 3.1 Å from the Tyr68 aromatic ring (**Fig. 3**F), which is consistent with a rare thiol-aromatic bond (*43*). However, this peak was weaker or absent in protomer B and the P2_1_/C2 structures (**Fig. S5**). There was a similar difficulty locating the Cys143 thiol deuterium atom: a positive peak in the F_O_-F_C_ NSL density map was present in protomer B of the P4_3_ structure, but absent in the other structures. These discrepancies may be due to incomplete HDX because the cysteines are buried, and/or the different resolutions of the neutron structures.

### Protein flexibility and hydrogen bond networks in Mac1

To better understand the structural basis for ADPr recognition in Mac1, we examined hydrogen bond networks in the functionally important loops that surround the active site. First, we analyzed variation across Mac1 crystal structures by aligning structures determined at cryogenic and room temperature from several different crystal forms (**Fig. 4**A/B). The resulting ensemble shows that structure variation is mainly restricted to two regions around the active site: the loop at that positions Phe156 in the adenosine binding site (the Phe156 loop), and the two loops that position Phe132 in the catalytic site (the Ile131 loop and the Lys102 loop). Next, we compared structural variation across structures determined at cryogenic temperature and ambient temperature (**Fig. 4**C). The Cα RMSF was highly correlated (Pearson r = 0.93), indicating that structural variation was independent of data collection temperature. The Cα B-factors were also highly correlated between the two temperatures (Pearson r = 0.92) (**Fig. 4**C). The region with the highest Cα B-factors at both temperatures was the Lys102 loop, which is consistent with the structural variation seen across different crystal forms (**Fig. 4**B).

**Fig. 4.**
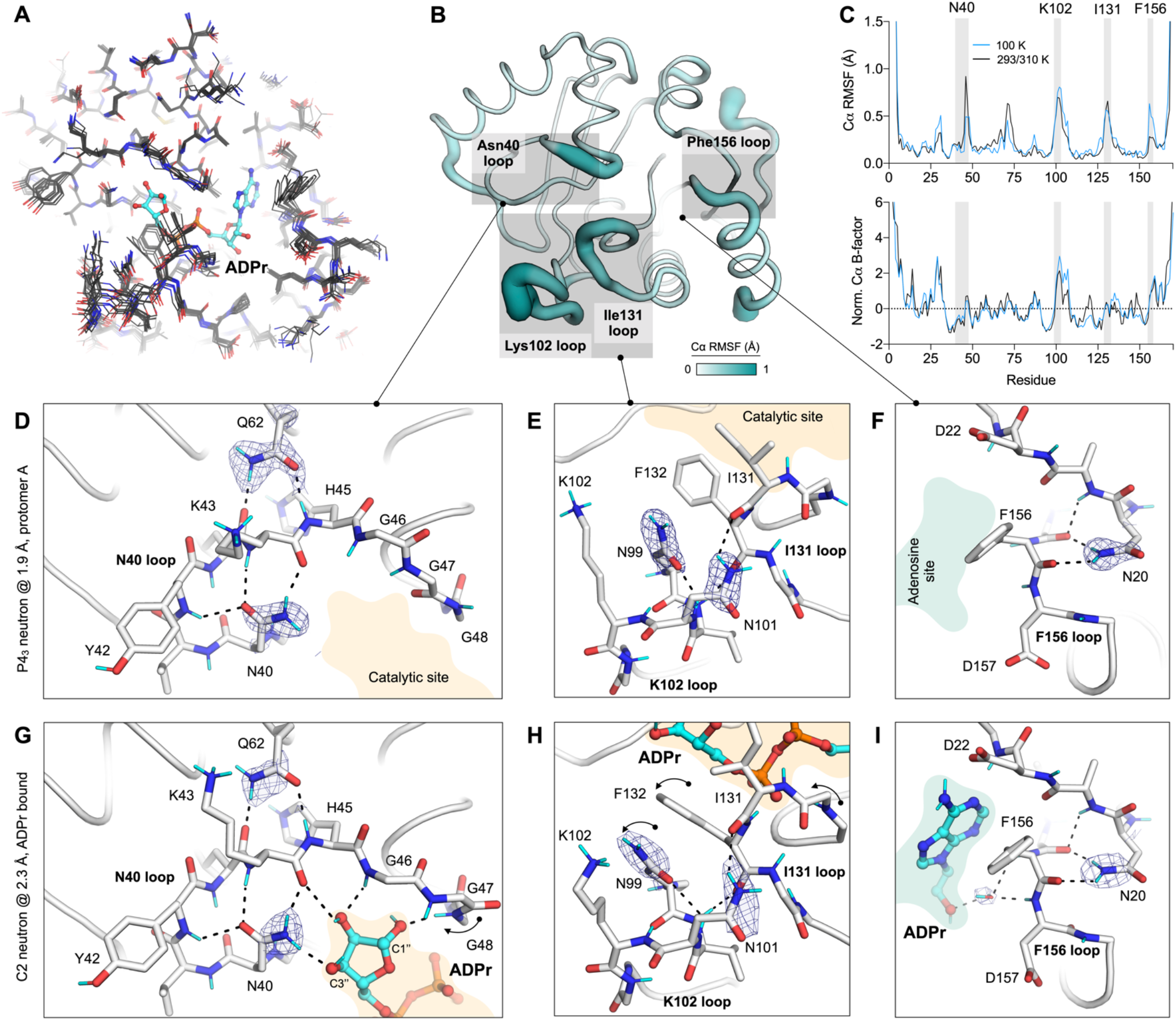
Protein flexibility and hydrogen bond networks in the Mac1 active site. (**A**) Alignment of P4_3_, P2_1_ and C2 Mac1 neutron and X-ray crystal structures determined at 100/293/310 K (PDB codes 7KQO, 7KQP, 7KR0, 7KR1, 7TWH, 7TX3, 7TX4, 7TX5). (**B**) Cα RMSF calculated from structures shown in (**A**) mapped onto the Mac1 structure. (**C**) Top: plot showing Cα RMSF of Mac1 structures determined at 100 K (black line) and those at 293/310 K (blue line). Bottom: plot showing Cα B-factors from Mac1 structure. B-factors were normalized by Z-score as described in the Methods section. (**D**,**E**,**F**) Active site hydrogen bond networks assigned based on NSL density maps for the P4_3_ neutron structure (protomer A, PDB code 7TX3). The protein is shown with a white stick/cartoon representation and the 2mF_O_-DF_C_ NSL density map is shown with a blue mesh (contoured at 2 σ around the asparagine/glutamine side chains). Hydrogen bonds (<3.5 Å) are shown with dashed black lines. (**G, H, I**) Same as (**D**,**E**,**F**), but showing the hydrogen bond networks in the C2 ADPr co-crystal structure (PDB code 7TX5). The NSL density maps are shown with a blue mesh (contoured at 3, 2 and 2.5 σ in **G, H** and **I**).

#### Asn40 network

The need for flexibility to allow substrate binding is often balanced by the need to stabilize a substrate for catalysis (*44, 45*). In Mac1, Asn40 plays an important role in substrate stabilization, based on its absolute conservation across evolutionarily diverse macrodomains, and loss of macrodomain function when Asn40 is mutated to alanine (*46*). In the ADPr-bound Mac1 neutron structure, Asn40 forms a hydrogen bond with the C3’’ hydroxyl of ADPr, with the side chain orientation clearly defined in the NSL density map (**Fig. 4**G). The Asn40 side chain is preorganized to make this interaction, based on the similarity in the side chain conformation in the apo and ADPr-bound structures (**Fig. 4**D/G, **Fig. S1**). This preorganization is achieved by hydrogen bonds to the backbone of Tyr42 and Lys44, which is in turn stabilized by hydrogen bonds to the side chain of Glu62 (**Fig. 4**D/G). This network is conserved at 100 and 293 K, across different Mac1 crystal forms and between the SARS-CoV and MERS macrodomains (*16*). This extensive network stabilizes Asn40, while allowing the adjacent glycine-rich loop (Gly46-48) to be flexible during ADPr binding.

#### Ile131/Lys102 network

Next, we examined hydrogen bond networks in the Ile131/Lys102 loops, which undergo a series of conformational changes upon ADPr binding (**Fig. 1**C). The orientation of Asn99 is clearly defined in NSL density maps in the three neutron structures (**Fig. 4**E/H). The side chain nitrogen is oriented towards the catalytic site, while the side chain oxygen is hydrogen bonded to the backbone nitrogen of Asn101. In the P4_3_ structure, the nearby Asn101 forms a hydrogen bond with the backbone of Ile131, connecting the two loops. This network is disrupted in the P2_1_ crystal form, where Asn101 forms a hydrogen bond with a symmetry mate. Overall, this network is conserved between the 100 K and 293 K P4_3_ X-ray structures, with only a slight shift in the Lys102 loop (**Fig. S6**). However, there is substantial variation in this loop in the ensemble of Mac1 determined using different crystal forms, and this variation matches the conformational change that occurs upon ADPr binding (**Fig. S6**). The Lys102 and Ile131 loop movement is correlated, with inter-loop hydrogen bonds maintained across the ensemble. In the SARS-CoV macrodomain, an arginine and aspartic acid are present at the position equivalent to Asn99 and Asn101. Although this removes a hydrogen bond between the Lys102 loop and the Ile131 loop, it introduces a new salt bridge to constrain the Asp98 side chain (equivalent to Asn99 in Mac1). Taken together, analysis of the Ile131/Lys102 loops shows how a key flexible region of Mac1 is supported by inter-loop hydrogen bond networks. This flexibility may be important for accommodating a wide range of ADP-ribosylated substrates.

#### Phe156 network

The adenosine portion of ADPr is recognised by hydrogen bonds to Asp22 and Ile23, and a water mediated contact with the Phe156/Asp157 backbone (**Fig. 1**C). The Phe156 loop is stabilized by two hydrogen bonds that connect the Asn20 side-chain to the Val155/Phe156 backbone (**Fig. 4**F/I). The orientation of the Asn20 side-chain is clearly defined in the NSL density maps obtained from both the P4_3_ and C2 crystals (**Fig. 4**F/I). Although multiple conformations of the Phe156 side-chain were seen in the cryogenic P4_3_ structure, only a single conformation was modeled in the 293 K X-ray structure (**Fig. S6**). However, in the Mac1 ensemble obtained from different crystal forms, the Phe156 side-chain is structurally diverse, matching the diversity seen for fragment-bound structures (*14*). Both SARS-CoV and MERS have an asparagine at the position equivalent to Phe156 and a cysteine at the position equivalent to Asn20 (**Fig. S6**). This indicates a degree of tolerance of the residues at this position, given that all three enzymes have similar ADPr binding properties (*9*). One consequence of this amino acid change is the potential for π-π stacking interactions at this site. Although the adenine portion of ADPr is too far to form π-π stacking interactions, we previously observed several fragments making close contacts with the Phe156 side-chain (*14*). An inhibitor that relies too heavily on stacking interactions with Phe156 may therefore have limited use as a broad-spectrum antiviral against coronaviruses.

### Water networks in Mac1

Water networks play an important role in protein-ligand recognition (*47, 48*), however their contribution to ligand binding is often difficult to quantify. Here, the combination of ultra-high resolution cryogenic X-ray structures (0.77 and 0.85 Å), a high-resolution room temperature structure (1.1 Å), structures determined using different crystal forms (P4_3_, P2_1_ and C2), high resolution neutron structures (1.9 Å and 2.3 Å), and more than 200 fragment bound structures offers an exceptional opportunity to study the role of water in macrodomain structure, function and ligand binding.

#### Active site water networks at room temperature

To examine water structure within the Mac1 active site, we first mapped water networks in the 1.1 Å room temperature X-ray structure determined using P4_3_ crystals (**Fig. 5**A/B). There were 19 active site water molecules in total: 14 in the catalytic site and five in the adenosine site. The catalytic site network encircles the Phe132 side chain, and is larger and more interconnected compared to the adenosine site network (**Fig. 5**A). To assess the tolerance of water networks to crystal packing, we compared the networks between the two protomers in the P4_3_ ASU (**Fig. 5**C). Although the adenine site of protomer B is occupied by the N-terminal residues of a symmetry mate, the water positions and B-factors were similar (RMSD = 0.40 Å, Pearson r for B-factors = 0.42). Next, we compared the water network from the 1.9/2.3 Å X-ray/neutron structure determined using the P2_1_ crystal. The water network was similar (RMSD = 0.45 Å), with only W10 and W16 not resolved in the P2_1_ structure. Together, these results indicate that the Mac1 active site water network is conserved across different crystal packing environments.

**Fig. 5.**
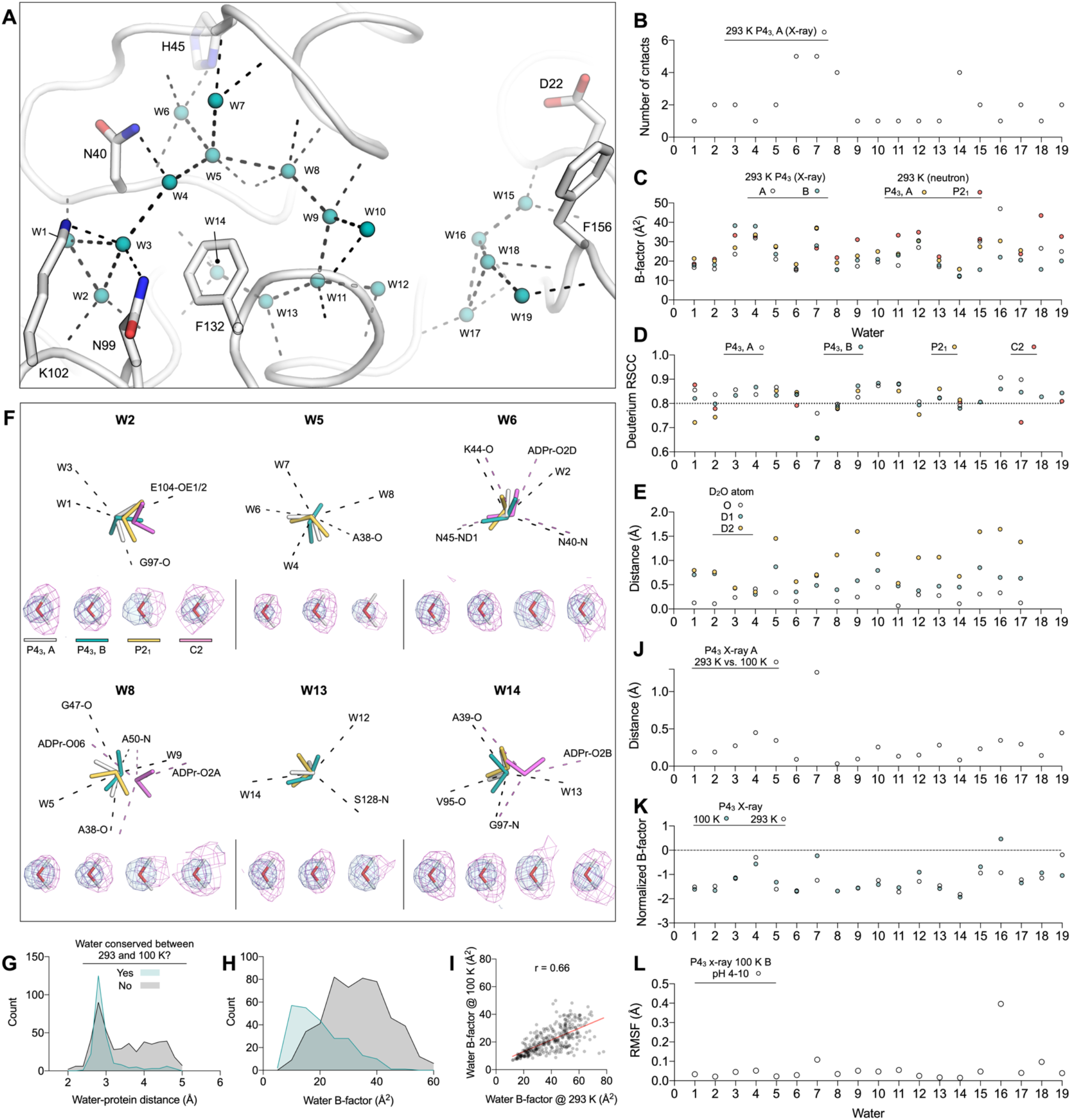
Mac1 active site water networks are robust to changes in crystal packing, temperature and pH. (**A**) Water network in the Mac1 active site from the 1.1 Å P4_3_ X-ray structure obtained at room temperature (PDB code 7TWH). Waters were considered part of the active site network if they were within 3.5 Å of an active site hydrogen bond acceptor/donor. For clarity, only the side chains of selected residues are shown. The protein is shown with white cartoon/sticks, the waters are shown as teal spheres, and hydrogen bonds with dashed black lines. (**B**) Plot showing the number of contacts for water molecules shown in (**A**). (**C**) Plot showing B-factors for the active site H_2_O/D_2_O molecules in the room temperature P4_3_ X-ray structure and the P2_1_ and C2 neutron structures. Solvent molecules are numbered according to (**A**). (**D**) Real-space correlation coefficients (RSCC) for active site D_2_O molecules in the P4_3_, P2_1_ and C2 neutron structures calculated using the 2mF_O_-DF_C_ NSL density map. A line is drawn at an RSCC=0.8, which has previously been used as a threshold for assessing whether a D_2_O is correctly oriented (*49*). (**E**) Variation in water orientations across 100 independent rounds of refinement. The plot shows distances between the average deuterium position of protomer A and B of the P4_3_ structure. For comparison, the D_2_O oxygen distances between the A and B protomers are shown. (**F**) Selected active site D_2_O molecules showing the 2mF_O_-DF_C_ NSL density map (purple mesh contoured at 2 σ) and 2mF_O_-DF_C_ electron density map (blue mesh/surface contoured at 2 σ). Hydrogen bonds are shown with dashed black lines. For the C2 structure, ADPr specific hydrogen bonds are shown with pink dashed lines. (**G**) Histogram showing water protein-distances in the 0.85 Å P4_3_ X-ray structure determined at 100 K (PDB code 7KQO). Waters were grouped based on whether a matching water molecule within 0.5 Å was found in the P4_3_ structure determined at 293 K (PDB code 7TWH). The histogram was generated with a bin width of 0.2 Å. (**H**) Same as (**G**) but showing B-factors for all water molecules in the 100 K P4_3_ structure. The histogram was generated with a bin width of 5 Å^2^. (**I**) Scatter plot showing correlation between water B-factor at 100 and 293 K. The red line shows a linear fit of the data using GraphPad Prism. (**J**) Plot showing distances between H_2_O oxygen atoms between protomer A of the P4_3_ X-ray structures determined at 293 and 100 K (PDB codes 7TWH and 7KQO). (**K**) Plot showing the B-factors of active site H_2_O molecules in the P4_3_ X-ray structures determined at 293 and 100 K. B-factors were normalized by Z-score using the B-factors from all the H_2_O molecules in a structure. (**L**) Plot showing the RMSF of active site water molecules calculated across the seven structures determined from pH 4 to 10 using the P4_3_ crystal form at 100 K. Because of pH-dependent binding of buffer components in the active site of protomer A, only the waters in protomer B are shown.

#### D_2_O orientations from neutron diffraction

Although the high resolution Mac1 X-ray crystal structures allow the accurate placement of water molecule oxygens, hydrogens are not visible at this resolution, which means that the orientation of water molecules is unclear. Neutron crystallography yields a more detailed map of water networks by determining the orientation of D_2_O molecules. To model D_2_O molecules in the neutron structures, deuterium atoms were added to D_2_O oxygens and their orientations were refined using the 2F_O_-F_C_ NSL density maps (*35*). Deuterium atoms were removed from D_2_O molecules that lacked density. Although all D_2_O molecules in the active site of the P4_3_ structure were modeled with deuterium, the lower resolution of the P2_1_ structure meant that six D_2_O molecules were modeled without deuterium (**Fig. S7**). The quality of water modeling was assessed by calculating the real-space correlation coefficient (RSCC) (*49*) (**Fig. 5**D). The RSCC values were consistently high, indicating that there was good agreement between D_2_O orientations and NSL maps. Next, we assessed the reproducibility of D_2_O orientation in refinement by running 100 independent rounds of refinement starting from coordinates where the D_2_O deuterium atoms were randomly shifted by 0.5 Å (**Fig. S8**). Deuterium atoms were included for all D_2_O molecules in this procedure. The RMSF of deuterium atoms calculated across the 100 structures provides a measure of the reproducibility of D_2_O orientations. For D_2_O molecules in the P4_3_ structure, RMSF values were low, consistent with highly reproducible D_2_O orientations (**Fig. S8**). RMSF values for W3, W4 W9, W15 and W18 were comparatively high in the P2_1_ structure, supporting their omission from the final model. To measure the agreement between equivalent D_2_O molecules across the two P4_3_ protomers, we calculated the distance between the average deuterium coordinates from the 100 structures (**Fig. 5**E). The D1/D2 positions for W3, W4, W6, W11 and W14 were all within 0.6 Å on average, consistent with conserved D_2_O orientations between the P4_3_ protomers. Based on B-factor and coordination number, W6 and W14 are the most ordered waters in the active site, and this matches their conserved orientation across the three Mac1 monomers (**Fig. 5**F).

#### Water networks in Mac1 are conserved from 100 K to 293 K

It is well established that crystal cryocooling causes crystal lattice contraction and remodeling of water networks at crystal packing interfaces (*50*–*52*). In the P4_3_ crystal form of Mac1, lattice contraction upon cryocooling was modest, with a 2% decrease in unit cell volume from 293 K to 100 K (PDB codes 7TWH versus 7QKO). To investigate the effect of cryocooling on Mac1 solvation, we compared the overall water structure in the 1.1 Å P4_3_ structure with the previously published 0.85 Å structure determined at 100 K (PDB code 7KQO). Although there were substantially fewer waters modeled in the room temperature structure relative to the cryogenic structure (360 versus 745), the positions and B-factors of the waters were highly conserved (**Fig. 5**G-I). Of the 360 waters modeled in the room temperature structure, 240 had a matching water within 0.5 Å in the cryogenic structure. Waters that were not conserved tended to be second-shell waters (i.e. they were more than 3.5 Å from protein atoms) and also tended to have higher B-factors. The resolution gap between the two structures (0.25 Å) is not large enough to explain the difference in the number of waters modeled (*53*). In line with previous observations (*52*), it seems likely that this difference is due to increased solvent disorder at room temperature. However, unlike previous observations (*52*), the apparent increase in solvent disorder is not coupled to increased protein disorder, based on the highly correlated disorder in the structures determined at cryogenic and room temperature (**Fig. 4**C, **Fig. S3**). Next, we compared active site water networks at 293 and 100 K (**Fig. 5**I/J). Water positions and B-factors were highly conserved (RMSD = 0.38, Pearson r for B-factors = 0.62). Unsurprisingly, the positions of waters with lower B-factors tended to be more similar.

#### Mac1 active site water networks are conserved from pH 4-10

Based on the similarity of the Mac1 structures determined using crystals grown at pH 6.5 and 9.5, Mac1 tolerates a broad pH range. To extend this analysis to a wider pH range and higher resolution, whilst eliminating the impact of different crystal packing, we performed a pH-shift experiment with the P4_3_ crystals. Briefly, crystals grown at pH 9.5 were soaked for four hours in a buffer composed of citric acid, HEPES and CHES, with pH values from 3 to 10 (*54*). There was no diffraction from crystals soaked at pH 3, despite the crystals appearing undamaged. From pH 4 to 10, crystals diffracted to better than 0.8 Å (data were truncated to 0.9 Å to achieve ∼100% completeness) (**Table S1**). Based on diffraction data recorded from at least three crystals at each pH, there was a small but significant pH-dependent change in until cell lengths (**Fig. S9**). This was most pronounced for the c-axis, which contracted 0.8% from pH 4 to 10. The unit cell changes may be linked to changes in the protonation states of His91 and His119 (**Fig. S10**). These residues are located at the interface between the two protomers in the P4_3_ ASU, and there was a pH dependent shift in the side chain χ_1_ and χ_2_ dihedral angles (**Fig. S10**). Surprisingly, we observed multiple buffer components binding to the oxyanion subsite (**Fig. S1**). Binding was pH dependent, with density for citrate between pH 4-6, weak density for acetate at pH 7 and 8, and weak density for CHES at pH 9 and 10. There was also weak density for CHES binding in an alternative site at pH 5 and 6. Although acetate was not added to the soaking buffer, it is a known contaminant of HEPES buffer (*55*). Binding of the buffer components in the oxyanion subsite led to substantial conformational changes in the Phe156 loop (**Fig. S9**), mirroring the changes seen across different crystal forms (**Fig. 4**A) and upon fragment binding (*14*). There was an increase in backbone disorder from pH 4 to 10 for the α-helices composed of residues 20-31 and 156-169, as well as the Lys102 loop (**Fig. S9**). Surprisingly, we observed a low occupancy alternative conformation of the Ile131 loop in protomer B at pH 4. This alternative conformation was associated with an alternative water network and shared some similarity to the ADPr-bound conformation (**Fig. S1**), although there was no flip in Ala129. Overall, the active site water network was highly similar across the entire pH range (**Fig. 5**L, **Fig. S9**).

#### Re-organization of water networks upon ADPr binding

To investigate the role of water in ADPr binding, we compared the active site water networks in the previously published apo and ADPr-bound structures determined using P4_3_ crystals (PDB code 7KQO and 7KQP) (**Fig. 6**A/B). In total, 11 waters are displaced by ADPr binding, either through direct displacement by ADPr (W4-5, W9-13, W15-16, W18) or through displacement by a shift in protein conformation (W3, displaced by Phe132). Of the remaining waters, six form bridging hydrogen bonds between ADPr and Mac1 (W6, W7, W8, W14, W17, W19). These waters tend to have more protein contacts and lower B-factors compared to the displaced waters (**Fig. 5**B/C). A notable exception is W19, which connects the oxyanion subsite to the C2’ hydroxyl and is stabilized in the ADPr-bound structure (**Fig. 6**A/C). There were minor adjustments in the bridging waters that bind ADPr: W6, W7, W8, W14 and W17 all shift by between 0.3-1 Å (**Fig. 6**B). In addition to the water displacement and reorganization, two new waters are found in the ADPr-bound structure. W_OXY_ extends the hydrogen bonding network near the oxyanion subsite, and W_PHOS_ is hydrogen bonded to the terminal ribose and adjacent phosphate of ADPr (**Fig. 6**A). In total, the re-organization of water networks upon ADPr binding fits a model where weakly bound waters (as measured by number of protein contacts and B-factors) are displaced, while tightly bound waters are recruited to form bridging ligand-protein interactions.

**Fig. 6.**
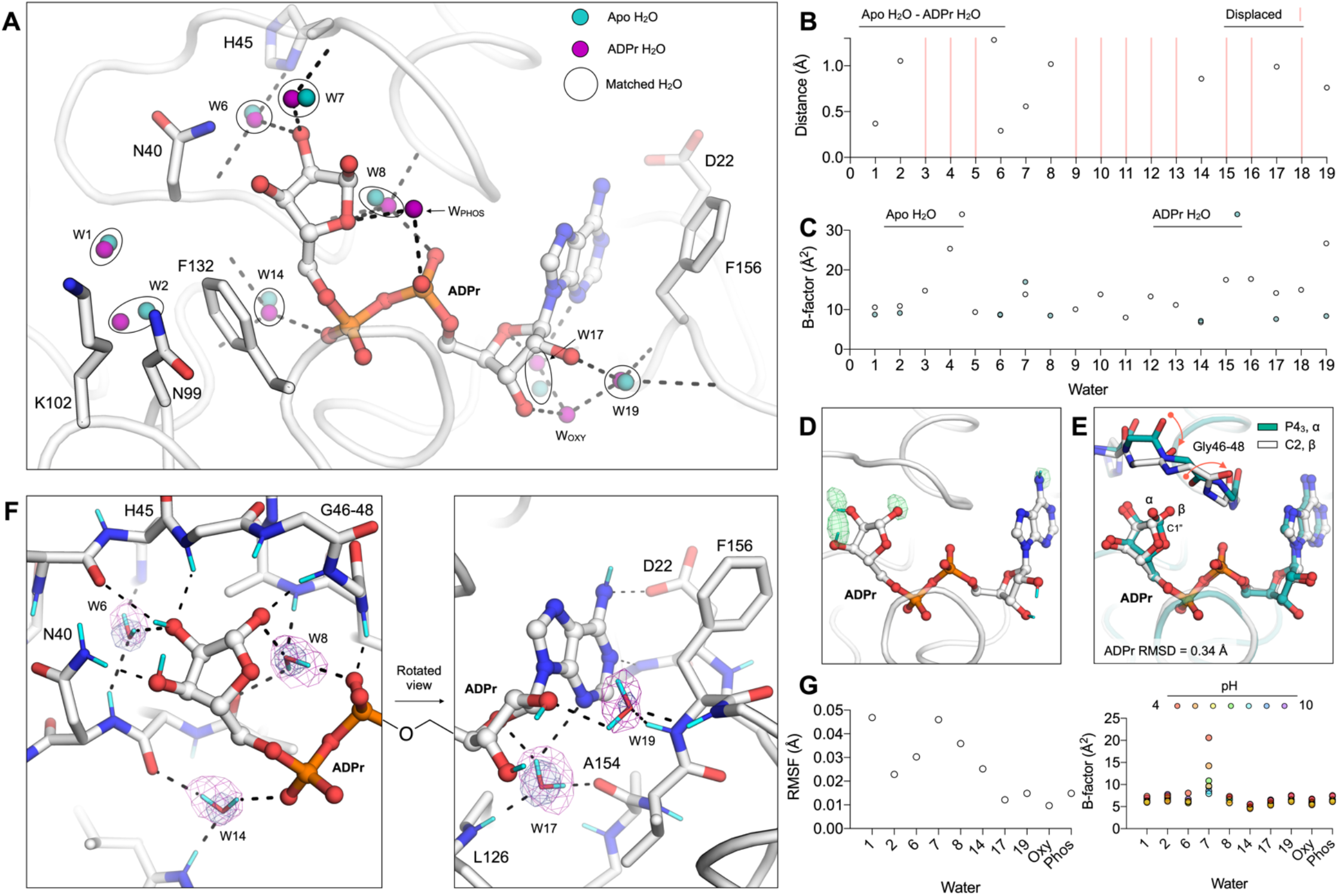
Reorganization of water networks upon ADPr binding. (**A**) Active site water positions from the structure of ADPr-bound Mac1 determined at 100 K (P4_3_ crystal, PDB code 7KQP) compared to the apo structure (PDB code 7KQO). For clarity, only selected side chains of the ADPr-bound structure are shown (white sticks). Hydrogen bonds are shown as dashed black lines. (**B**) Plot showing distances between the water molecules shown in (**A**). (**C**) Plot showing the B-factors for waters shown in (**A**). (**D**) Evidence for hydrogen-deuterium exchange in ADPr co-crystallized with Mac1 (C2, PDB code 7TX5). The mF_O_-DF_C_ NSL density map calculated after joint neutron/X-ray refinement but prior to adding deuterium atoms to ADPr is shown with green mesh (contoured at +3 σ). No density was observed for the C2’ and C3’ hydroxyl deuteriums. (**E**) Alignment of the ADPr-bound Mac1 structures determined using P4_3_ and C2 crystals. The two configurations of the terminal ribose are marked (α and β), and the conformational change required to bind the α configuration in the C2 crystal is shown with red arrows. (**F**) Hydrogen bond networks in the ADPr-bound Mac1 structure determined using neutron diffraction (C2, PDB code 7TX5). The 2mF_O_-DF_C_ NSL density map is shown for bridging water molecules with a purple mesh (contoured at 2.5 σ for W6, W8, W14 and W17, and at 2 σ for W19). The 2mF_O_-DF_C_ electron density map is contoured at 2 σ (blue mesh). Deuterium atoms are colored cyan and hydrogen bonds are shown with dashed black lines. (**G**) Left: plot showing the RMSF of active site water molecules calculated across the seven ADPr-bound structures determined from pH 4 to 10. Right: B-factors for water molecules in the ADPr-bound structures from pH 4 to 10.

Next, we examined hydrogen bond networks in the Mac1-ADPr in the C2 co-crystal system using neutron diffraction. Although the ADPr used for co-crystallization was hydrogenated, HDX occurred when the crystal was soaked in the deuterated buffer. This is supported by positive peaks in the F_O_-F_C_ NSL density map calculated prior to modeling ADPr deuterium atoms (**Fig. 6**D). In the neutron structure, the terminal ribose adopts the β-configuration, with rearrangement of the Gly46-48 loop accommodating the C1’’ hydroxyl (**Fig. 6**E). This configuration contrasts with the P4_3_ structure, where crystal packing interactions preclude rearrangement of the Gly46-48 loop, which forces the terminal ribose to adopt the α-configuration (**Fig. 6**E) (*14*). Despite the different configurations of the terminal ribose, and rearrangement of the Gly46-48 loop, the ADPr binding pose was very similar between the C2 and P4_3_ structures (RMSD = 0.34 Å, excluding the C1’’ hydroxyl). Likewise, the positions of bridging D_2_O molecules were highly conserved (RMSD for W6, W8, W14, W17 = 0.46 Å). Although there was a 1.2 Å shift in the position of W19, the hydrogen bonding network with the oxyanion subsite is maintained. Of the ADPr-specific bridging water molecules (W_OXY_ and W_PHOS_), only W_OXY_ was identified in the C2 structure. The absence of W_PHOS_ is due to crystal packing interactions: a close contact between Ala69 of a symmetry mate forces the Ile131 side chain to rotate and occupy the W_PHOS_ position (**Fig. S1**).

Refinement of the positions of ADPr deuteriums, and the orientations of bridging D_2_O molecules, allows the hydrogen bond networks that underpin ADPr binding to be examined (**Fig. 6**F). In the catalytic site, W8 forms hydrogen bonds with the C1’’ hydroxyl, the backbone oxygen of Ala38, the backbone nitrogen of Ala50 and one of the ADPr phosphate oxygens. This highly coordinated D_2_O is appropriately oriented to act as a water nucleophile in the putative substrate-assisted hydrolysis mechanism (*16*), with the D_2_O oxygen oriented towards the C1’’ carbon. Also in the catalytic site, W6 forms a bridging interaction between the deprotonated Nδ1 of His45 and the C2’’ hydroxyl of ADPr. The nearby Asn40, which plays a key role in Mac1 function (*16*), forms a 3 Å hydrogen bond with the C3’’ hydroxyl. In the adenosine site, W19 acts as a bridge between the oxyanion subsite and the C2’ hydroxyl (**Fig. 6**F). The second bridging water in this site, W17, forms hydrogen bonds to the backbone oxygen of Ala154, the backbone nitrogen of Leu126, and is within hydrogen bonding distance of both the N3 nitrogen and the O4’ oxygen of ADPr. In summary, the structure of ADPr bound to Mac1 determined by neutron diffraction has revealed the precise configuration of hydrogen bond networks in the active site, and highlights the role of bridging water molecules in ADPr binding.

#### ADPr binding is conserved from pH 4-10

The high resolution X-ray structures used to investigate changes in water networks upon ADPr binding were obtained using crystals grown at pH 9.5 (*14*). To investigate ADPr binding across a broader pH range, we repeated the pH-shift experiment, this time soaking crystals in a buffer supplemented with 20 mM ADPr. Similarly to the apo pH-shift experiment, the diffraction properties of the crystals at pH 4-10 were excellent, with reflections recorded beyond 0.8 Å at all pHs. Data were truncated to 0.9 Å to achieve 100% completeness, with the exception of the dataset at pH 9, which was recorded at 17 keV resulting in ∼100% completeness at 0.84 Å (**Table S1**). There was clear density for ADPr in the active site at all pHs, with the refined occupancy exceeding 95% (**Fig. S1**). Like the apo crystals, we observed a pH-dependent contraction along the c-axis of the unit cell, although the a/b-axis expansion was absent (**Fig. S9**). Compared to the apo structures, there was only a slight increase in disorder going from pH 4 to 10, consistent with stabilization of protomer A by ADPr binding. At pH 4, there was weak density for an alternative conformation of His45 in protomer A (**Fig. S10**). The conformational change involved an 8 Å shift in the His45 side chain, with an adjustment in the backbone torsion angles of Gly46/47 and rearrangement of water molecules to occupy the cavity left by His45 (**Fig. S10**). The change in His45 confirmation is likely induced by protonation at pH 4, indicating that the doubly protonated tautomer is sterically incompatible with the buried position. Despite the conformational change in this active site loop, the coordinates and B-factors of ADPr were highly similar across the entire pH range (**Fig. S9**), as were the positions and B-factors of active site water molecules (**Fig. 6**G). These pH-shift experiments show that Mac1 can bind ADPr from pH 4 to 10 in the P4_3_ crystals.

#### Role of water networks in fragment binding

To investigate the role of bridging water molecules in the binding of non-native ligands to Mac1, we analyzed water networks in the previously published structures of 232 fragments bound to Mac1 (*14*). The fragment structures were determined as part of a large-scale screen using X-ray crystallography, and included 196 fragments bound in the Mac1 active site. Our previous analysis revealed that in some cases, fragment binding occurred through extensive water-mediated interactions (*14*). Here, we extend this analysis to include the newly identified water networks from the room temperature X-ray and neutron crystal structures. First, we created a map of bridging water density in the adenosine site using multi-Gaussian spreading (*56*) (**Fig. 7**A). Waters were classified as bridging if they were within 3.5 Å of both a fragment hydrogen bond donor/acceptor and a protein hydrogen bond donor/acceptor. Of the 196 fragments binding in the Mac1 active site, 130 fragments had at least one bridging water, with an average of 2.1 bridging waters per fragment. To quantify the extent of clustering around apo waters in the active site, we measured the distance between bridging waters and the nearest apo water (**Fig. 7**B). Clusters with at least 10 waters were found near W10/W12, and W15-19. The largest cluster was near W17, with 64 waters within 2 Å and 42 waters within 0.2 Å. W17 sits at the base of the adenine subsite, and is involved in bridging interactions with ADPr (**Fig. 6**F). Out of the 64 fragments that hydrogen bond to W17, the fragment atom was an oxygen hydrogen bond donor in 59 of them. The remaining five contained either nitrogen hydrogen bond donors or acceptors. Visualization of the fragment atoms that hydrogen bond to W17 shows that this interaction is highly flexible (**Fig. 7**D). Hydrogen bonds between W17 and fragment atoms were distributed from 2.6-3.5 Å, and over a wide range of angles (**Fig. 7**E-G). The flexibility in bridging interactions with W17 may explain the large number of carboxylic acid containing fragments that bound to the oxyanion subsite (*14*). Of the 64 fragments interacting with W17, 55 were carboxylic acids. In particular, W17 allows carboxylic acids to interact with the oxyanion subsite in several distinct ways, based on bond connectivity and carboxylic acid orientation (**Fig. 7**H). Bridging waters were also identified beyond the adenosine site, with three nearby clusters visible in the water density map (labeled WB1, WB2 and WB3 in **Fig. 7**A).

**Fig. 7.**
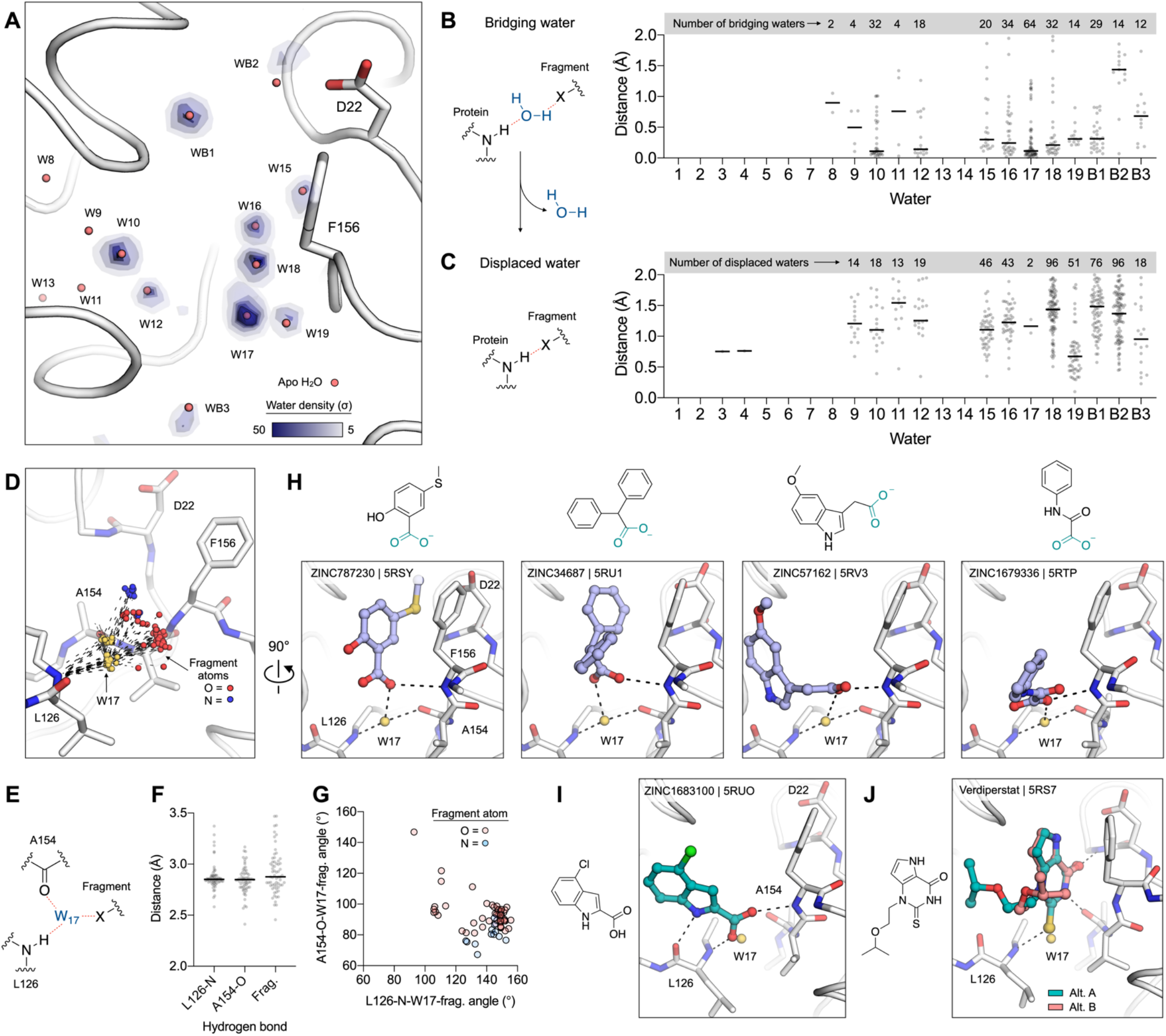
Water networks mediate fragment binding in the adenosine site of Mac1. **(A**) Structure of Mac1 showing density of fragment-protein bridging water molecules, calculated using the 232 previously reported fragment structures (*14*). The P4_3_ 100 K structure (PDB code 7KQO) is shown with a white stick/cartoon representation. Water density was calculated with the GROMAPS tool (*56*) and is contoured from 5 to 50 σ. (**B**) Left: chemical structure showing an example of a bridging water molecule. Right: distances from bridging waters to apo waters calculated for all fragment structures. (**C**) Left: chemical structure showing an example of water displacement. Right: the minimum fragment-water molecule for each fragment structure. (**D**) W17 acts as a bridge to 64 fragments binding in the adenosine site. The Mac1 structure (PDB code 7KQO) is shown, and W17 from the 64 fragments is shown with yellow spheres. The hydrogen bonds to Leu126 and Ala154 are shown with black dashed lines, and the fragment atoms are shown with spheres colored by atom type. (**E**,**F**,**G**) The W17-Leu126/Ala154 hydrogen bonds are conserved across the 64 bridging fragments, whereas the W17-fragment bonds are variable, based on distance (**F**) and angle (**G**). (**H**) W17 mediates diverse interactions with carboxylic-acid containing fragments and the oxyanion subsite. The fragments are shown with blue sticks. (**I**,**J**) W17 is displaced by 2 out of the 178 fragments binding in the adenosine site. Two conformations of Verdiperstat were observed.

Next, we examined the displacement of waters from the adenosine network by fragment binding. A water was classified as displaced if a fragment atom was within 2 Å of the water in the apo structure (**Fig. 7**C). On average, each fragment binding in the Mac1 active site displaced 1.6 waters from the active site network. The most frequently displaced waters were W15, W16, W18 and W19, which matches the large number of fragments bound in the adenine and oxyanion subsites (*14*). Waters that were highly ordered in the apo structure were displaced infrequently or not at all. Notably, W17 was only displaced by two fragments out of 178 binding in the adenosine site (**Fig. 7**C). In both cases, the fragments form extensive hydrogen bond networks to neighboring residues (**Fig. 7**I/J). The indole-containing ZINC1683100 forms hydrogen bonds to the backbone nitrogen/oxygen of Leu129 and to the oxyanion subsite (**Fig. 7**I), while the thioketone of Verdiperstat forms hydrogen bonds with the Leu126 backbone nitrogen and the adenine subsite (**Fig. 7**J). In summary, the perturbation of water networks in the adenosine site by fragment binding is similar to the perturbation seen upon ADPr binding (**Fig. 6**A): waters that are highly ordered in the apo structure are exploited to form bridging interactions in the ligand-bound enzyme. Furthermore, analysis of the fragment-bound structures shows that the majority of waters in the adenosine site network are available for bridging interactions or displacement (**Fig. 7**B/C). This analysis highlights the potential for solvent networks to be exploited in the design of Mac1 ligands, through the introduction of groups that extend bridging hydrogen bond networks or through the targeted displacement of water molecules.

## Discussion

The neutron structures reported here advance our understanding of Mac1 structure and function in three main ways. First, the structures allowed the protonation state of histidine residues to be assigned at pH 6.5 and 9.5 in the apo and ADPr-bound enzyme (**Fig. 3**C). Second, the neutron structures helped to identify hydrogen bond networks that control functionally important loop flexibility in and around the active site (**Fig. 4**D-I). Third, the neutron structures enabled the orientation of ordered water molecules to be determined, helping to construct a detailed map of water networks in the Mac1 active site (**Fig. 5, Fig. 6**). These three main findings collectively inform the potential for inhibitor optimization and determination of the catalytic mechanism of Mac1.

The combination of diffraction data collection using different crystal forms, at different temperatures, and at different pHs show that the water networks in the Mac1 active site are surprisingly conserved across a range of perturbations. In contrast, ADPr binding leads to reorganization of the water networks with the displacement of 11 waters, the stabilization of two new waters and small shifts in the remaining waters (**Fig. 6**A). The displacement of weakly bound waters, and the recruitment of tightly bound waters in bridging interactions is consistent with previous large scale surveys of ligand-water-protein interactions (*57*–*59*). Although ADPr recognition involves a series of changes in Mac1 conformation that either introduce new polar contacts with ADPr (e.g. the Gly48/Ala129 peptide flips) or remove steric blocks (e.g. the coupled Phe132/Asn99 side chain rotation), small shifts in tightly bound waters are seen in the Mac1-ADPr complex (**Fig. 6**B). This may reflect a general principle governing water-mediated ligand-protein contacts: small shifts in the positions of water molecules leads to ligand stabilization that would be difficult to achieve through direct protein-ligand contacts. Our analysis of bridging water molecules in the structures of ∼200 fragments bound in the adenosine site suggests that these principles extend to non-native ligands (**Fig. 7**B). Notably, W17 acts as a bridging water in the ADPr complex, and also forms bridging interactions with 64 fragments.

The detailed map of active site water molecules presented here will help ligand discovery efforts against Mac1 for three main reasons. First, the water network analysis shows that bridging waters in native and non-native ligands are clustered around the position of water molecules in the apo structure (**Fig. 6**A, **Fig. 7**A). This correlation can be leveraged by minimizing the energy penalty associated with solvent reorganization. The involvement of the same waters in bridging interactions with a large collection of chemically diverse fragments is promising for virtual screening efforts against Mac1, because it shows that water positions can be taken directly from the apo structure, which is consistent with previous observations (*60*). Second, the water network analysis identified buried waters that can be targeted for displacement. This strategy is frequently invoked in ligand optimization efforts (*48, 61*–*63*), but it is unclear how broadly applicable this strategy is for optimizing potency. The large number of bridging waters identified here provide an unmatched opportunity to systematically examine the thermodynamics of water displacement (for example, displacement of W17 by the indole-containing ZINC1683100 in **Fig. 7**i). Third, receptor models for virtual screening can be improved by using the experimentally determined water orientations from the Mac1 neutron structures (**Fig. 5**F, **Fig. 6**F). The orientation of water molecules included in receptor models can have a major impact on the success of virtual screening (*64*). One caveat to using the neutron-determined orientations is that only orientations of highly ordered water molecules could be determined with high confidence (**Fig. 5**D/E). Fortunately, these waters are most frequently involved in bridging interactions. Simulation-based methods might yield an improved model of the orientations of disordered water molecules (*65*).

We found that the conformational ensemble of Mac1 was unusually robust to temperature and pH perturbation in the crystal. Room temperature crystallography can reveal low occupancy, functionally relevant conformational states of proteins (*51, 66*–*68*). The 1.1 Å room temperature structure of Mac1 reported here, in combination with the previously reported 0.85 Å structure determined using the same crystal form at 100 K (*14*), allowed a detailed analysis of temperature dependent changes in structure and function. Overall, the structure and flexibility at 100 and 293 K were remarkably similar, indicating that unlike several recently reported examples (*68*–*70*), Mac1 displays minimal temperature dependent structural changes *in crystallo* (**Fig. 4**C). Similarly, assuming that the crystals have equilibrated to the target pH in the ∼4 hour soaks conducted here (*71*), the pH shift experiments revealed that the Mac1 active site residues and water networks are remarkably robust (**Fig. 6**I/J), and this is matched by the invariance in ADPr binding across the entire pH range (**Fig. 6**G/H). One caveat to the pH-shift experiments is the pH-dependent binding of citrate, acetate and CHES in the oxyanion subsite (**Fig. S1**). Although this complicated the analysis of pH-dependent structural changes, this interference would apply equally to solution-based experiments, and highlights the protein binding potential of commonly used buffers (*72*). A valuable use of the pH-shift experiment described here would be to investigate the pH-dependent binding of ionizable inhibitors to Mac1. Collectively, these experiments highlight a conspicuous problem related to using crystallography to study protein flexibility: the same tightly packed crystal lattices that help to deliver the high resolution information needed to identify subtle changes in structure may lead to protein stabilization that masks the same structural changes.

We discovered conformational diversity by comparing Mac1 structures across different crystal forms (**Fig. 4**C). Importantly, this ensemble of structures captures most of the structural diversity seen upon ADPr and fragment binding (**Fig. S6**). One notable exception is the Phe132 side chain, which adopts a similar conformation in all of the apo crystal structures, but is stabilized in a different rotamer in the ADPr-bound structure (**Fig. S6**). Surprisingly, the relatively constant position of Phe132 across the apo structures occurs despite substantial variation in the Asn99/Lys102 loops (**Fig. S6**). This stability may help to explain why our fragment screening campaign failed to identify any fragments binding in the terminal ribose site (*14*): fragments are unable to overcome the energetic penalty required to stabilize the Phe132 side-chain in the ADPr-bound “flipped” conformation. The lack of fragments in this site may also be due to the stability and the associated enthalpic cost of displacing the highly ordered waters in this site (e.g. W13 and W14). While our previous virtual screening campaign focussed solely on identifying fragments binding to the adenosine site, a worthwhile extension would be to use the ADPr-bound conformation as a receptor model to target the catalytic site.

Although the physiologically relevant substrates of Mac1 are unknown, macrodomains are thought to primarily hydrolyse ADPr conjugated to glutamate or aspartate side chains at the C1’’ position (**Fig. 1**A) (*73*). Based on X-ray and neutron crystal structures of the SARS-CoV-2 macrodomain with ADPr bound, two substrate binding modes, distinguishable based on the conformation of the terminal ribose (**Fig. 8**A/B) and leading to different potential catalytic mechanisms are plausible. In the first mode, which matches the conformation seen in the P4_3_ crystal structure, the terminal ribose adopts the α configuration, with the conjugated peptide exiting the active site through the gap between the Gly46-48 loop and the Ile131 loop. Although this gap is narrow in the apo structures, it widens by ∼1 Å in the ADPr-bound structure, and we previously identified a fragment wedged between these loops (PDB code 5RVI) (*14*). In this binding mode, the proximity of the C1’’ carbon to the phosphate groups, and the absence of another suitable general base, implicates the previously proposed substrate-assisted hydrolysis mechanism with either W_PHOS_ or W8 acting as a nucleophile (*15, 16*) (**Fig. 8**C). If the hydrolysis reaction involves an S_N_2 type nucleophilic substitution at the C1’’ carbon, then we favor W8 as the water nucleophile, because it can approach the sp^3^ carbon with a collinear trajectory relative to the peptide leaving group (**Fig. 8**C) and the neutron crystal structure with ADPr-bound shows that the water oxygen is oriented towards the C1’’ carbon (**Fig. 6**G). This is in contrast to the water nucleophile proposed by Jankevicius *et al*. (*15*) in hMacroD2, which is equivalent to W_PHOS_ in Mac1 (**Fig. 8**E).

**Fig. 8.**
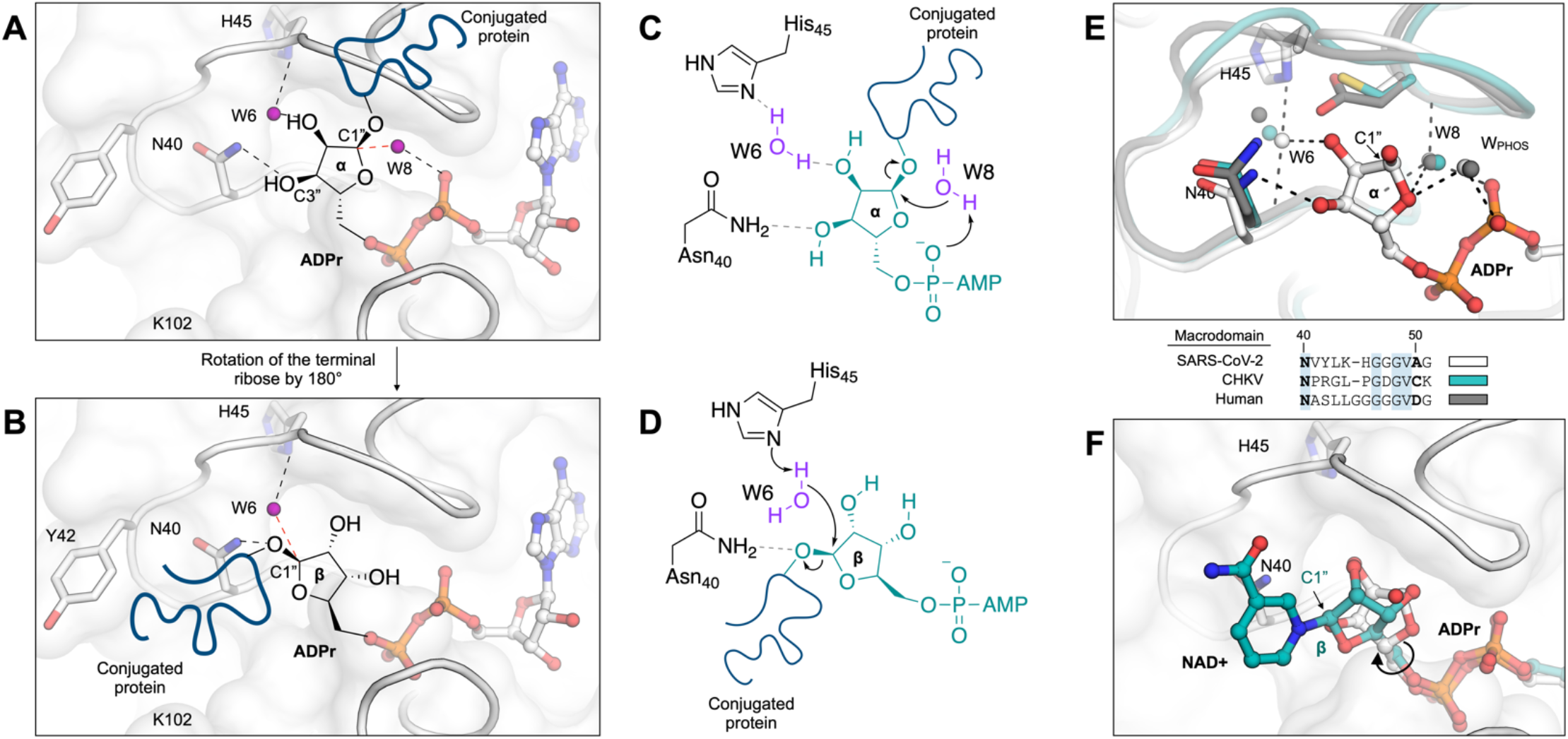
Mechanism of ADPr-ribose hydrolysis catalyzed by the SARS-CoV-2 NSP3 macrodomain. (**A**) Composite image showing the ADPr-bound Mac1 structure (white sticks/cartoon/surface, PDB 7KQP). ADPr is shown in the configuration that is compatible with a substrate-assisted mechanism. The terminal ribose adopts the α-configuration and W8 acts as the water nucleophile. For clarity, only the side chains of selected residues are shown. Hydrogen bonds are shown with dashed black lines and the W6-C1’’ trajectory is shown with a dashed red line. (**B**) Same as (**A**), but showing the β-configuration of the terminal ribose that is compatible with His45 acting as a general base to activate W6 as a nucleophile. (**C, D**) Chemical structures showing the two possible mechanisms for ADPr hydrolysis. (**E**) Top: structural alignment of the SARS-CoV-2 macrodomain (PDB code 7KQP), Chikungunya virus macrodomain (PDB code 3GPO) and the human macrodomain hMacroD2 (PDB code 4IQY), all in complex with ADPr. Bottom: protein sequence alignment of residues equivalent to residues 40-51 from the SARS-CoV-2 macrodomain. (**F**) X-ray crystal structure of the NSP3 macrodomain from the Tylonycteris bat coronavirus HKU4 in complex with NAD+ (PDB code 6MEB). The terminal ribose is rotated ∼180° relative to ADPr. This configuration matches the model proposed in (**B**/**D**).

In the second possible ADPr binding mode, the ribose adopts the β configuration and the terminal ribose is rotated 180° about the C4”-C5” bond relative to the configuration observed in the P4_3_ crystals (**Fig. 8**B). In this binding mode, a substrate-assisted mechanism is unlikely because the C1’’ carbon would be >6 Å from the diphosphate. However, several lines of evidence support the nearby His45 as a general base in the hydrolysis mechanism for coronaviral macrodomains (**Fig. 8**D). First, the neutron structures show that His45 has the appropriate orientation and protonation state to act as a general base, with a deprotonated Nδ1 nitrogen and a D_2_O molecule hydrogen bonded between His45 and ADPr (**Fig. 3**D). Second, His45 is conserved across all known coronavirus macrodomains, with mutation to alanine eliminating ADPr-hydrolysis activity (*16*). It should be noted that His45 is absent in alphavirus macrodomains and the human macrodomain hMacroD2, although both macrodomains contain a nearby nucleophilic residue (Asp102 in hMacroD2 and Cys34 in the Chikungunya virus macrodomain) (**Fig. 8**E). Third, the binding mode with the terminal ribose rotated ∼180° was observed in the structure of the macrodomain from Tylonycteris bat coronavirus HKU4 with NAD+ bound (*74*) (**Fig. 8**F). The nicotinamide group is oriented towards a groove formed between Tyr42 and Lys102. We previously identified several fragments binding in this site (*14*), and have uncovered conformational flexibility consistent with the site being able to accommodate a range of peptide substrates exiting the Mac1 active site (**Fig. S6**). Moreover, docking of the potential Mac1 substrate PARP-1 into the Mac1 active site places a portion of the PARP-1 peptide in this groove (*74*). Finally, this binding mode hints at an alternative role for Asn40 in ADPr hydrolysis. Asn40 is strictly conserved across macrodomains and mutation to alanine eliminates activity (*10*). In the substrate-assisted mechanism, this functional importance has been attributed to the hydrogen bond formed between Asn40 and the C3’’ hydroxyl of ADPr. In the alternative mechanism proposed here (**Fig. 8**B/D), Asn40 could have a direct role by stabilizing the peptide leaving group. Discriminating between the two ADPr binding modes would require a structure of Mac1 in complex with a suitable substrate mimic (e.g. a non-hydrolyzable ADP-ribosylated protein or peptide). As has been suggested previously (*16*), these two possible mechanisms raise the intriguing possibility that distinct mechanisms of ADPr-hydrolysis have evolved from highly similar protein scaffolds.

In summary, we have determined neutron crystal structures of the SARS-CoV-2 NSP3 macrodomain using multiple crystal forms across the apo and ADPr-bound states. We assigned the protonation states of histidine residues, and mapped hydrogen bond networks that control the flexibility of active site loops that are involved in the recognition of ADPr. A comprehensive analysis of water networks in the Mac1 active revealed that the water networks are conserved across different crystal forms, between 100 and 293 K and from pH 4 to 10. The water networks reorganize upon ADPr binding, and play a key role in the molecular recognition of non-native ligands in the adenosine site. The series of structures presented here increase our understanding of macrodomain catalytic mechanisms and will serve as a resource to guide the design of new Mac1 inhibitors.

## Materials and Methods

### P4_3_ crystals

#### Protein expression and purification

The gene encoding SARS-CoV-2 NSP3 Mac1 (residues 3-169) was cloned into a pET-22b(+) expression plasmid with a TEV-cleavable N-terminal 6-His tag (Genscript). The protein was expressed and purified as described previously (*14*).

#### Crystallization for neutron diffraction

Neutron diffraction quality crystals were grown using sitting drop vapor diffusion. Crystallization drops were set up on 22 mm siliconized glass coverslips (Hampton Research, HR3-231) with 65 μl protein, 60 μl reservoir (34% PEG 3000, 100 mM CHES pH_aparent_ 9.1 (pD 9.5)) and 5 μl seed stock. Seed stocks were prepared as described previously (*14*) and were stored at -80°C. The sitting drops were equilibrated against a 4 ml reservoir solution stored in a small plastic beaker, using two 10 cm plastic petri dishes taped together to create a sealed environment. Crystals reached their maximum size of approximately 2×2×0.5 mm after one week. To increase the deuterium content, crystals were transferred from crystallization drops into an artificial reservoir solution containing 34% PEG 3000 and 100 mM CHES pH 9.5, prepared using D_2_O. Because crystals were tightly bound to the coverslips, it was necessary to use a fine needle to dislodge them. Crystals were allowed to soak in the artificial reservoir solution for 24 hours, before being transferred into a 2 mm thin walled quartz capillary (Hampton Research, HR6-150). To keep crystals hydrated, 20 μl of the D_2_O-containing reservoir solution was pipetted into one end, and the capillaries were sealed with wax at both ends (Hampton Research, HR4-328).

#### Neutron/X-ray diffraction data collection, reduction and refinement

Neutron time-of-flight diffraction data were collected at room temperature on the MaNDi instrument at the SNS (*32, 75*). An incident neutron wavelength bandpass of 2-4 Å was used with a Δϕ of 10° between frames. A total of 10 diffraction images were collected with an average exposure of 16 h per frame. Exposure was set to 80 Coulomb of charge on the SNS target. At 1.4 MW operation, charge accumulates at a rate of approximately 5 C/h. Fluctuation in exposure time occurs because the accelerator does not operate 100% of the time. The neutron dataset was reduced using the Mantid package (*76*) and integrated using three dimensional profile fitting (*77*). The data were wavelength normalized using LAUENORM from the LAUEGEN suite (*78*–*80*).

Following neutron diffraction data collection, an X-ray dataset was collected on the same crystal at room temperature using a microfocus rotating anode X-ray diffractometer (Rigaku HighFlux HomeLab equipped with a MicroMax-007 HF X-ray generator, Osmic VariMax optics and an Eiger R 4M hybrid photon counting detector). A total of 360 diffraction images were collected with a Δϕ of 0.5° and an exposure time of 10 s per frame. X-ray diffraction data were integrated using the CrysAlis Pro software suite (Rigaku Inc.). Diffraction data were then reduced and scaled using the Aimless (*81*) program from the CCP4 suite (*82*). The X-ray data collection statistics are shown in Table S1.

The Mac1 structure was initially refined against the X-ray data only. Phases were obtained by molecular replacement with Phaser (*83*) using the coordinates of the apo Mac1 as the search model (monomer A of PDB code 7KQO, stripped of alternative conformations and waters). To debias refinement from the coordinates used for molecular replacement, the B-factors of 7KQO were reset to 20 Å^2^ and the coordinates were randomly offset by 0.3 Å using phenix.pdbtools. The structure was refined with phenix.refine (*84*) using default parameters and five macrocycles. Manual model building was performed with Coot (*85*). In the early stages of refinement, waters were added automatically in phenix.refine to peaks in the mF_O_-DF_C_ map 3.5 σ or higher. CHES was modeled into the adenosine site with restraints generated using phenix.elbow (*86*). The occupancy of CHES was set to 50% initially and was refined automatically in subsequent steps. Positive peaks in the mF_O_-DF_C_ electron density map that overlapped with CHES were modeled as partially occupied water molecules. The occupancy of CHES and the partially occupied waters was constrained to 100% in phenix.refine, with the final CHES occupancy equal to 62%. In the later stages of refinement, hydrogens were added to the model using Reduce (*42*) run through phenix.ready_set. Hydrogens were refined at their riding positions. Cycles of refinement and model building continued until *R*_free_ no longer decreased. The coordinates were prepared for joint neutron/X-ray refinement by adding deuterium atoms to all exchangeable positions and D_2_O molecules with phenix.ready_set. Therefore, exchangeable positions were modeled with both a hydrogen and deuterium using A/B alternate location identifiers, and the deuterium occupancy was automatically refined. Joint refinement was performed using phenix.refine (*35*) with hydrogens/deuteriums refined individually. Hydrogens/deuterium ADPs were refined and neutron distances were used for hydrogen/deuterium bonds. After five macrocycles of refinement, the 2mF_O_-DF_C_ (unfilled) and the mF_O_-DF_C_ NSL density maps were inspected in Coot and modifications made to the coordinates. To assign histidine protonation states, mF_O_-DF_C_ NSL density maps were inspected after refinement with either the doubly protonated or doubly deprotonated forms (**Fig. S5**). Water orientations were automatically refined with phenix.refine. The fit of D_2_O molecules into 2mF_O_-DF_C_ NSL density maps was assessed by visual inspection, and the orientations of D_2_O molecules were adjusted when necessary. Deuteriums were deleted from water molecules that lacked supporting density.

#### Crystallization for high resolution room temperature X-ray diffraction

Crystals were grown using hanging drop vapor diffusion in 24-well crystallization plates (Hampton, HR3-171). Crystallization drops contained 4 μl Mac1 at 40 mg/ml, 2 μl seeds and 2 μl reservoir consisting of 32% PEG 3000 and 100 mM CHES pH 9.5 with 500 μl reservoir in each well. Crystals reached a maximum size of approximately 1×1×0.25 mm after four days. Crystals were looped using dual thickness MicroLoops (Mitegen, M5-L18SP) mounted on B3-3 magnetic bases (Mitegen, GB-B3-R). MicroRT tubing (Mitegen, RT-T1) with 5 μl of reservoir in the end was placed over the crystals to prevent dehydration, and the tubing was sealed to the crystal mount with epoxy.

#### High resolution room temperature X-ray diffraction data collection, reduction and refinement

X-ray diffraction data was collected at the Stanford Synchrotron Radiation Lightsource (SSRL) using beamline 12-1. The cryojet temperature was set to 293 K and crystals were mounted remotely from an SSRL crystallization plate (Mitegen, M-CP-111-095). To assess the susceptibility of crystals to radiation damage, datasets were collected from four similarly sized crystals with the X-ray dose increased by adjusting the beam transmittance while keeping the other data collection parameters constant (see **Table S1**). The absorbed dose for each dataset was calculated with RADDOSE-3D (*87*) using the parameters listed in **Table S1**. X-ray diffraction data was indexed, integrated and scaling used XDS (*88*) and merged with Aimless (*81*). Data collection statistics are shown in **Table S1**. Phases were obtained by molecular replacement with Phaser using the same search model as the lower resolution X-ray experiment. Refinement was performed with phenix.refine using default settings and five macrocycles at each step. Water molecules were added automatically in phenix.refine using the same settings as the lower resolution X-ray refinement. ADPs were refined isotropically in the first two cycles of refinement and anisotropically in the remaining cycles. After three cycles of refinement and model building, hydrogens were added to the model using Reduce. Hydrogens were refined using a riding model, with their ADPs refined isotropically. In the final two stages of refinement, the weights for the stereochemical and ADP restraints were relaxed (using the wxc_scale=1.5 and wxu_scale=1.5 flags in phenix.refine). Refinement and model building continued until *R*_free_ no longer decreased. In the final refinement cycle, automatic water modeling was switched off, and waters were placed manually into peaks in the mF_O_-DF_C_ electron density map. Coordinates and structure factor intensities were deposited in the PDB with accession codes 7TWF (73 kGy), 7TWG (153 kGy), 7TWH (290 kGy) and 7TWI (539 kGy).

#### pH-shift experiment

Crystals for the pH shift experiment were grown at 20°C using 96-well sitting drop crystallization plates (Hampton Research, HR3-125). Crystallization drops contained 200 nl protein, 100 nl reservoir solution (28% PEG 3000, 100 mM CHES pH 9.5) and 100 nl seed stock, with 30 μl in the reservoirs. Crystals grew to their maximum size in two days with dimensions approximately 0.4×0.4×0.1 mm. Crystals were transferred from crystallization plates into a buffer containing 22 mM citric acid, 33 mM HEPES and 44 mM CHES (2:3:4 molar ratio of the buffer components) (*54*) and 28% PEG 3000, with the pH adjusted from 3 to 10. Crystals were soaked for four hours, then vitrified in liquid nitrogen. The soaks with ADPr were performed in the same manner, but the buffers were supplemented with 20 mM ADPr (Sigma, A0752) (from a 500 mM stock prepared in H_2_O). Diffraction data were collected at beamline 8.3.1 of the Advanced Light Source (ALS) for 3-6 crystals at each pH. Data collection strategy and statistics are shown in **Table S1** for the best diffracting crystals at each pH. Phases were obtained by molecular replacement using the same search model as the room temperature experiments. Refinement was performed with phenix.refine using a similar protocol to the high resolution room temperature structures. ADPs were refined anisotropically (excluding hydrogens) and the occupancy of all water molecules was refined. Hydrogens were refined using a riding model, and in the final two stages of refinement the weights for stereochemical and ADP restraints were relaxed (using the wxc_scale=2 and wxu_scale=2 flags in phenix.refine). pH dependent binding of buffer components were observed in the apo crystal structures, these were refined with restraints generated by phenix.elbow for CHES or from the internal CCP4 library for citrate and acetate. ADPr binding was confirmed by inspection of mF_O_-DF_C_ electron density maps, with restraints generated by phenix.elbow. Coordinates and structure factor intensities were deposited in the PDB with accession codes 7TWJ (pH 4, apo), 7TWN (pH 5, apo), 7TWO (pH 6, apo), 7TWP (pH 7, apo), 7TWR (pH 8, apo), 7TWQ (pH 9, apo), 7TWS (pH 10, apo), 7TWT (pH 4, ADPr), 7TWV (pH 5, ADPr), 7TWW (pH 6, ADPr), 7TWX (pH 7, ADPr), 7TWY (pH 8, ADPr), 7TX0 (pH 9, ADPr), 7TX1 (pH 10, ADPr). Data reduction and refinement statistics are presented in **Table S1**, with the statistics produced with phenix.table_one.

### P2_1_/C2 crystals

#### Protein expression and purification

The gene encoding SARS-CoV-2 2-170 NSP3 Mac1 was cloned into a pET-11a plasmid (Bio Basic) and transformed into *E. coli* (BL21-DE3). A detailed procedure for Mac1 expression and purification has been published elsewhere (*36*).

#### Crystallization for neutron diffraction

Purified Mac1 was stored at 4°C and filtered through a 0.2 μm centrifugal filter prior to crystallization trials. Initial crystals grown in a 20 μl drop of Mac1 (∼17.5 mg/ml) mixed with 28% PEG 4000, 0.1 M MES pH 6.5 at a 1:1 ratio at 16°C were used to prepare a microseed stock with seed beads (Hampton Research). Large-volume crystals of ligand-free Mac1 were grown using a Hampton nine-well sandwich box set up with 150 μl drops of Mac1 (∼19 mg/ml) mixed at a 1:1 ratio with 25% PEG 4000, 0.1 M MES pH 6.5 seeded with 0.2 μl microseeds (1:3000 dilution). Crystals were left to grow undisturbed at 16°C for >2 months resulting in a crystal measuring ∼1.9 mm at the longest edge. The crystal was mounted in a fused quartz capillary accompanied by a plug of 27% PEG 4000 prepared in 100% D_2_O. For the ADP-ribose complex crystal, Mac1 (∼15 mg/ml) was mixed with ADPr at a 1:2 molar ratio, incubated at room-temperature for 30 minutes. The crystallization conditions were identical to those for ligand-free Mac1 crystal growth. The crystal drops were seed struck with a cat whisker from a small Mac1+ADPr co-crystal previously grown in a 24-well plate using a 24% PEG 4000, 0.1 M MES pH 6.5 well solution, and then the nine-well plate was incubated at 16°C. A Mac1+ADPr crystal measuring ∼2.4 mm at the longest edge was mounted in quartz capillary with 27% PEG 4000 prepared in 100% D_2_O. At the time of mounting, both crystallization drops were measured with a microelectrode to be pH 6.5.

#### Neutron/X-ray diffraction data collection and reduction

The full neutron diffraction dataset for ligand-free Mac1 crystal at pH 6.5 was collected using the MaNDi instrument and the data were processed in a similar fashion as for the ligand-free Mac1 crystal at pH 9.5. Neutron diffraction data for the Mac1-ADPr complex were collected at the IMAGINE instrument at the High Flux Isotope Reactor (HFIR) at Oak Ridge National Laboratory (*37, 89, 90*). The crystal was held stationary at room temperature, and diffraction data for each image were collected for 23 hours using neutrons between 3.3-4.5 Å. During the experiment, the next 23-hour diffraction image was collected after crystal rotation by Δϕ = 10°. IMAGINE neutron diffraction data were processed using the Daresbury Laboratory LAUE suite program LAUEGEN modified to account for the cylindrical geometry of the detector (*79, 91*). The program LSCALE (*92*) was used to determine the wavelength−normalization curve using the intensities of symmetry-equivalent reflections measured at different wavelengths. No explicit absorption corrections were applied. These data were then merged in SCALA (*93*). The neutron data collection statistics are shown in **Table S1**. Following the neutron data collection, the room-temperature X-ray diffraction datasets were collected from the same crystals in the same manner as described previously for the P4_3_ crystals. The X-ray data collection statistics are shown in **Table S1**.

#### Joint neutron/X-ray refinement

Joint neutron/X-ray refinement was performed in the same manner as the P4_3_ neutron/X-ray experiment. Phases were obtained by molecular replacement using the same coordinates as the P4_3_ experiments. For the ADPr co-crystal structure, binding of ADPr was confirmed by inspection of mF_O_-DF_C_ electron density map. Restraints for the β isomer of ADPr were generated with phenix.elbow from the corresponding SMILES string. Coordinates and structure factor intensities were deposited in the PDB with accession code 7TX4 for the P2_1_ apo structure and 7TX5 for the C2 ADPr-bound structure.

### Structural analysis

#### Protein

RMSF values were calculated using a python script in PyMOL (version 2.5.1, Schrodinger). First, structures were aligned using Cα atoms. Second, average coordinates for each Cα were calculated across the structures. Third, the RMSF was calculated relative to the average Cα coordinates. Normalized Cα B-factors (B’) were calculated using the following equation: B’ = (B-μB)/σB, where B is the Cα B-factor, μB is the average B-factor across all Cα atoms and σB is the Cα B-factor standard deviation. For residues with alternative conformations, only the A conformation was included in the calculations of RMSF/average B-factor. To color structures by RMSF, the B-factor column in PDB files was replaced with the RMSF values. RMSD values were calculated using PyMOL with five cycles of outlier rejection.

The protonation states of histidine residues were assigned based on NSL density maps after refinement with either the doubly or deprotonated tautomers (maps are shown in **Fig. S5**). For the doubly protonated tautomers, the occupancy of the Nε2/Nδ1 deuterium atoms was set to 100%, and automatic refinement of occupancy was disabled.

#### Water

RSCC values for D_2_O molecules were calculated using phenix.real_space_correlation. The number of contacts made by water molecules was defined as the number of protein nitrogen or oxygen atoms within 3.5 Å of a water, with distances calculated using PyMOL. Normalized B-factors (B’) for water molecules were calculated with the following equation: B’ = (B-μB)/σB, where B is the water B-factor, μB is the average B-factor across all water molecules and σB is the water B-factor standard deviation.

To assess the reproducibility of water orientation refinement, we ran 100 independent rounds of refinement starting from the three jointly refined Mac1 structures. Deuterium atoms were added to all D_2_O oxygen atoms prior to refinement, and a random offset of 0.5 Å was applied to deuterium atoms using phenix.pdbtools. This procedure meant that each round of refinement started with a random distribution of D_2_O orientations, but with the same oxygen position. The variability in D_2_O orientations after refinement was quantified by calculating the RMSF value for each water deuterium atom across the 100 structures (**Fig. S8**). Agreement between the orientation of D_2_O molecules in the active sites of protomer A and B of the P4_3_ structure was assessed by calculating the distance between the average position of each D_2_O deuterium from the 100 refined structures (**Fig. 5**E).

The conservation of water molecules between structures was calculated with a python script in PyMOL. First, symmetry mates (including waters) within 5 Å of protein heavy atoms were generated and appended to each coordinate file. Next, the structures were aligned by Cα atoms, and the closest water in the structure being compared was determined, as well as the B-factors for both waters and the minimum distance to a protein hydrogen bond donor/acceptor.

The density map for bridging water molecules in the 232 previously reported Mac1 fragment-bound structures (*14*) was calculated using the GROMAPS tool (*56*) in GROMACS (*94*). First, water molecules related by symmetry within 5 Å of protein heavy atoms were appended to each fragment structure. Next, bridging water molecules were identified by selecting waters within 3.5 Å of both a protein and a fragment hydrogen bond acceptor/donor. No angle requirements were applied to the selection of bridging water molecules. Next, the fragment coordinates were aligned to protomer A of the 0.85 Å P4_3_ apo structure (PDB code 7KQO) and the coordinates of the bridging water molecules were saved and concatenated to form a single trajectory in PDB format. For GROMACS to accept the trajectory, it was necessary to add up to six waters to each structure so that the number of waters in each structure was the same. These waters were selected at least 25 Å from the active site. The grid resolution for calculating the density map was set to 0.5 Å.

## Acknowledgements

The SARS-CoV-2 Mac1 expression plasmid (pMCSG53) was provided by Dr. Andrzej Joachimiak (Argonne National Laboratory) with support from the National Institute of Allergy and Infectious Diseases, National Institutes of Health, Department of Health and Human Services, under Contract HHSN272201700060C. We thank Dr. Hugh M. O’Neill from ORNL for assistance during expression of the P2_1_/C2 constructs. Structural biology applications used in this project at UCSF were compiled and configured by SBGrid (*95*). This research was supported by the DOE Office of Science through the National Virtual Biotechnology Laboratory (NVBL), a consortium of DOE national laboratories focused on response to COVID-19, with funding provided by the Coronavirus CARES Act. This research used resources at the Spallation Neutron Source and the High Flux Isotope Reactor, which are DOE Office of Science User Facilities operated by the Oak Ridge National Laboratory. The Office of Biological and Environmental Research supported research at ORNL’s Center for Structural Molecular Biology (CSMB), a DOE Office of Science User Facility. This research used resources of the Spallation Neutron Source Second Target Station Project at Oak Ridge National Laboratory (ORNL). ORNL is managed by UT-Battelle LLC for DOE’s Office of Science, the single largest supporter of basic research in the physical sciences in the United States. The synchrotron X-ray diffraction data used to determine Mac1 structures from pH 4 to 10 were collected at beamline 8.3.1 of the Advanced Light Source. The ALS, a U.S. DOE Office of Science User Facility under Contract No. DE-AC02-05CH11231, is supported in part by the ALS-ENABLE program funded by the National Institutes of Health, National Institute of General Medical Sciences, grant P30 GM124169-01. The synchrotron X-ray diffraction data used to determine Mac1 structures at room were collected at beamline 12-1 of the Stanford Synchrotron Radiation Lightsource (SSRL). Use of the SSRL, SLAC National Accelerator Laboratory, is supported by the U.S. Department of Energy, Office of Science, Office of Basic Energy Sciences under Contract No. DE-AC02-76SF00515. The SSRL Structural Molecular Biology Program is supported by the DOE Office of Biological and Environmental Research, and by the National Institutes of Health, National Institute of General Medical Sciences (P30GM133894). L.C. acknowledges support by the NIH (R01-GM071939). This work was supported by NIH GM123159, NSF Rapid 2031205, and a TMC Award from the UCSF Program for Breakthrough Biomedical Research, funded in part by the Sandler Foundation (to J.S.F.).

## Author contributions

G. Correy expressed, purified and crystallized the P4_3_ construct, reduced X-ray diffraction data for P4_3_ crystals, refined and analyzed the neutron and X-ray structures, prepared the figures and wrote the manuscript. D. Kneller crystallized the P2_1_/C2 construct, collected and reduced neutron/X-ray data from the P2_1_/C2 construct. G. Phillips expressed and purified the P2_1_/C2 construct. S. Pant expressed and purified the P2_1_/C2 construct. S. Russi provided user support at SSRL, and assisted with the data collection strategy at SSRL. A. Cohen provided user support at SSRL, and assisted with the data collection strategy at SSRL. G. Meigs provided user support at ALS, and assisted with the data collection strategy at ALS. J. Holton provided user support at ALS, and assisted with the data collection strategy at ALS. S. Gahbauer edited the manuscript. M. Thompson assisted with the room temperature X-ray data collection strategy. A. Ashworth supervised work and arranged funding. L. Coates collected and reduced neutron/X-ray data for P2_1_/C2 construct. A. Kovalevsky collected and reduced X-ray and neutron data for the P2_1_/C2 construct and wrote the manuscript. F. Meilleur collected and reduced neutron/X-ray data for the P4_3_ construct and wrote the manuscript. J. Fraser supervised work, arranged for funding and wrote the manuscript.

## Data and materials availability

All data generated or analyzed during this study are included in this article and its Supplementary Information. Crystallographic coordinates and structure factors for all structures have been deposited in the Protein Data Bank with the following accessing codes: 7TWF, 7TWG, 7TWH, 7TWI, 7TWJ, 7TWN, 7TWO, 7TWP, 7TWQ, 7TWR, 7TWS, 7TWT, 7TWV, 7TWW, 7TWX, 7TWY, 7TX0, 7TX1, 7TX3, 7TX4, 7TX5.

