## Supplementary Information for "The mechanisms of catalysis and ligand binding for the SARS-CoV-2 NSP3 macrodomain from neutron and X-ray diffraction at room temperature"

### Supplementary Figures

**Supplementary Figure 1.** Binding of buffer components and ADP-ribose in the Mac1 active site.

**Supplementary Figure 2.** Summary of room temperature X-ray diffraction experiments using $P4_3$ Mac1 crystals.

**Supplementary Figure 3.** Comparison of backbone amide hydrogen-deuterium exchange across the three Mac1 neutron structures.

**Supplementary Figure 4.** Comparison of Mac1 structures determined at 293 K from $P4_3$, $P2_1$ and C2 crystals.

**Supplementary Figure 5.** Assignment of histidine and cysteine protonation states based on NSL / electron density maps.

**Supplementary Figure 6.** Flexibility and hydrogen bond networks in the Mac1 active site.

**Supplementary Figure 7.** Evidence for $D_2O$ orientations from neutron scattering length density maps.

**Supplementary Figure 8.** Assessing the variability in water orientations in joint X-ray/neutron refinement with phenix.refine.

**Supplementary Figure 9.** Investigating Mac1 conformational diversity with pH-shift crystallography.

**Supplementary Figure 10.** pH-induced conformational changes in Mac1 histidines.

### Supplementary Tables

**Supplementary Table 1.** Data collection and refinement statistics for crystal structures reported in this work (spreadsheet, Table_S1).

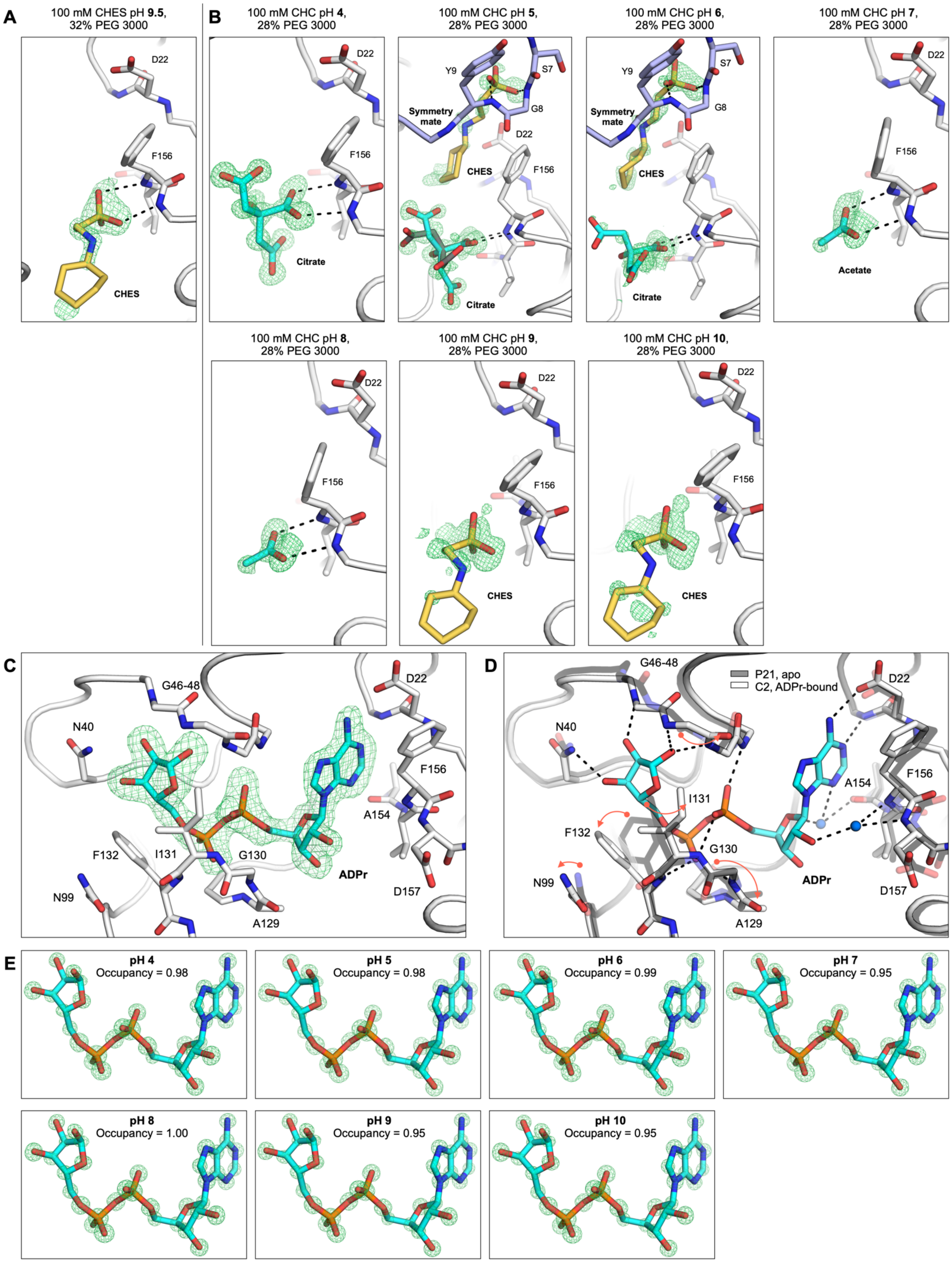
A
100 mM CHES pH 9.5, 32% PEG 3000
D22
F156
CHES
B
100 mM CHC pH 4, 28% PEG 3000
D22
F156
Citrate
100 mM CHC pH 5, 28% PEG 3000
Y9
S7
G8
Symmetry mate
D22
CHES
F156
Citrate
100 mM CHC pH 6, 28% PEG 3000
Y9
S7
G8
Symmetry mate
CHES
Citrate
100 mM CHC pH 7, 28% PEG 3000
D22
F156
Acetate
100 mM CHC pH 8, 28% PEG 3000
D22
F156
100 mM CHC pH 9, 28% PEG 3000
D22
F156
CHES
100 mM CHC pH 10, 28% PEG 3000
D22
F156
CHES
C
G46-48
N40
D22
F156
A154
F132
I131
G130
ADPr
N99
D157
A129
D
G46-48
P21, apo
C2, ADPr-bound
D22
N40
A154
F156
I131
F132
G130
ADPr
N99
D157
A129
E
pH 4
Occupancy = 0.98
pH 5
Occupancy = 0.98
pH 6
Occupancy = 0.99
pH 7
Occupancy = 0.95
pH 8
Occupancy = 1.00
pH 9
Occupancy = 0.95
pH 10
Occupancy = 0.95

**Fig. S1.** Binding of buffer components and ADP-ribose in the Mac1 active site. (**A**) CHES bound in the oxyanion subsite of protomer A of the $P4_3$ neutron structure (PDB code 7TX3). The $mF_O$-$DF_C$ electron density map calculated prior to modeling CHES is shown (green mesh contoured at +3 σ). The protein is shown with white sticks/cartoon and hydrogen bonds are shown with dashed-black lines. (**B**) pH dependent binding of buffer components in the adenosine site of Mac1 for the $P4_3$ crystals were soaked with 100 mM CHC buffer (22 mM citric acid, 33 mM HEPES and 44 mM CHES). The $mF_O$-$DF_C$ electron density map calculated prior to modeling the buffer components is shown with green mesh (contoured at +3 σ). Acetate was modeled at pH 7 and 8. Although it was not added to the CHC buffer, it is a known contaminant of HEPES (*55*). The positive $mF_O$-$DF_C$ electron density in the oxyanion subsite at pH 9 and 10 suggests that CHES is bound with low occupancy. However, the weak/diffuse nature of the density meant that CHES was not modeled. (**C**) ADPr bound in the Mac1 active site in the C2 X-ray/neutron structure. The $mF_O$-$DF_C$ electron density map prior to modeling ADPr is shown (green mesh contoured at +3 σ). (**D**) Structural alignment of the $P2_1$ and C2 neutron structures showing the conformational changes upon ADPr binding (red arrows). For clarity, only side chains of selected residues are shown. Key hydrogen bonds are shown with dashed black lines. (**E**) $mF_O$-$DF_C$ electron density maps calculated prior to ADPr modeling for the seven Mac1 crystal structures determined from pH 4 to 10 with the $P4_3$ crystals (green mesh contoured at 8 σ).

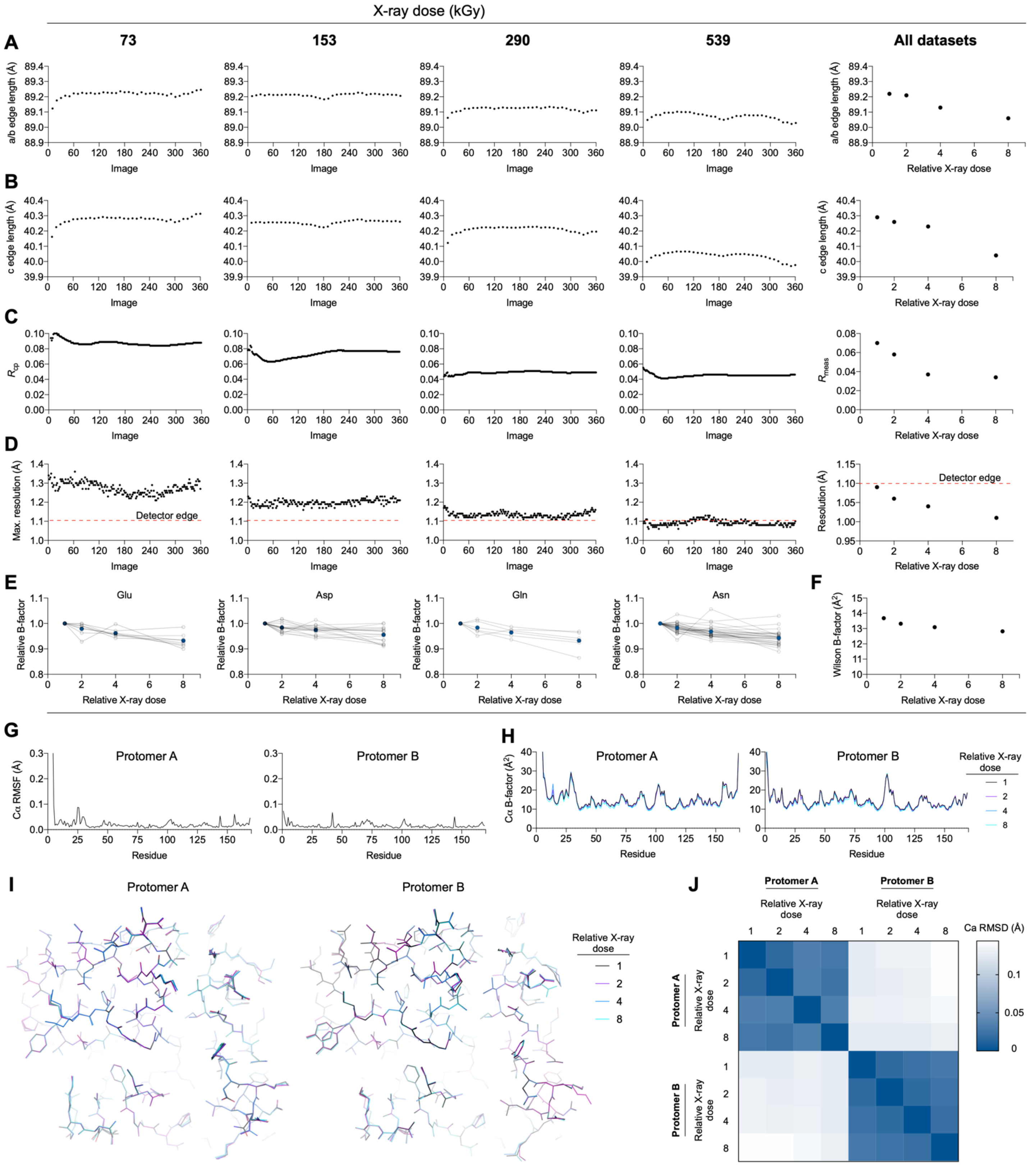


**Fig. S2.** Summary of room temperature X-ray diffraction experiments using P4$_3$ Mac1 crystals. (**A**) Plots showing unit cell edge lengths for the a/b axis refined by XDS for every 10 images (5° rotation). Each plot represents a dataset collected with increasing X-ray dose from four different, similarly sized crystals. (**B**) Same as (**A**) but showing the unit cell edge lengths for the c axis. (**C**) Plot showing the $R_{CP}$ statistic calculated by Aimless as a function of image (*96*). $R_{meas}$ is shown after truncating the data to 1.1 Å. (**D**) Plot showing the maximum resolution calculated by Aimless as a function of diffraction image (based on I/σ(I)>1). The overall dataset resolution is shown (based on $CC_{1/2}>0.3$) (*97*). All datasets were truncated to 1.1 Å to achieve ~100% completeness. (**E**) Plots showing the glutamate (Cδ, Oε1, Oε2), aspartate (Cγ, Oδ1, Oδ2), glutamine (Cδ, Oε1, Nε2) and asparagine (Cγ, Oδ1, Oδ2) B-factors at each X-ray dose relative to the lowest dose dataset. Residues with alternative conformations modeled were excluded. (**E**) Plot showing Wilson B-factors for each dataset. (**G**) Plot showing the Cα RMSF calculated across the Mac1 structures refined from the four datasets collected at 293 K. (**H**) Plot showing the Cα B-factors for the four 293 K Mac1 structures. (**I**) Structural alignment of the four 293 K Mac1 structures (PDB code 7TWF, 7TWG, 7TWH, 7TWI). (**J**) Heatmap showing the Cα RMSD between the four 293 K Mac1 structures.

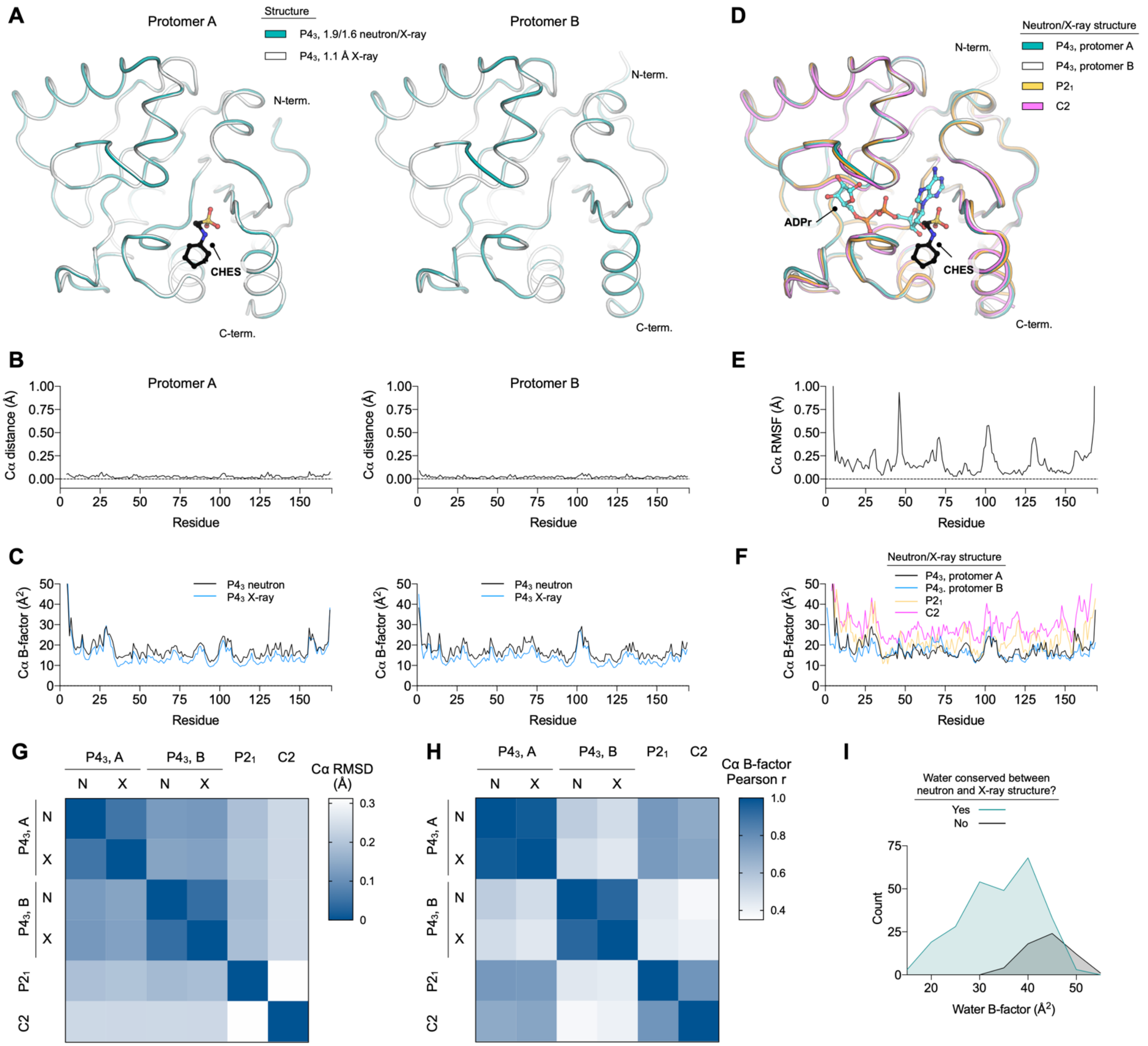


**Fig. S3.** Comparison of Mac1 structures determined at 293 K from P4$_3$, P2$_1$ and C2 crystals. (**A**) Alignment of the 1.9/1.6 Å neutron/X-ray structure (P4$_3$, PDB code 7TX3) with the 1.1 Å X-ray structure (P4$_3$, PDB code 7TWH). Alignment was performed separately for the A and B protomers. (**B**) Plot showing Cα distances calculated between the Mac1 structures shown in (**A**). (**B**) Plot showing Cα B-factors calculated for the Mac1 structures shown in (**A**). (**D**) Alignment of both protomers of the 1.9/1.6 Å neutron/X-ray structure (P4$_3$, PDB code 7TX3) with the 2.35/1.90 neutron/X-ray structure (P2$_1$, PDB code 7TX4) and the 2.30/1.95 neutron/X-ray structure of ADPr-bound Mac1 (C2, PDB code 7TX5). (**E**) Plot showing Cα RMSF values calculated across the four structures shown in (**D**). (**F**) Plot showing Cα B-factors for the four structures shown in (**D**). (**G**) Heatmap showing the Cα RMSD calculated between the 1.1 Å X-ray structure (P4$_3$, PDB code 7TWH) and the three neutron structures (P4$_3$, P2$_1$ and C2). In the heatmap legend, N denotes a neutron structure and X a X-ray structure. (**H**) Heat map showing the correlation of Cα B-factors of the structures shown in (**D**). (**I**) Histogram showing the B-factors for all water molecules in the 293 K P4$_3$ X-ray structure (PDB code 7TWH). Waters are grouped based on whether a matching water molecule was found within 0.5 Å in the P4$_3$ neutron structure (PDB code 7TX3). The histogram was generated with a bin width of 5 Å$^2$.

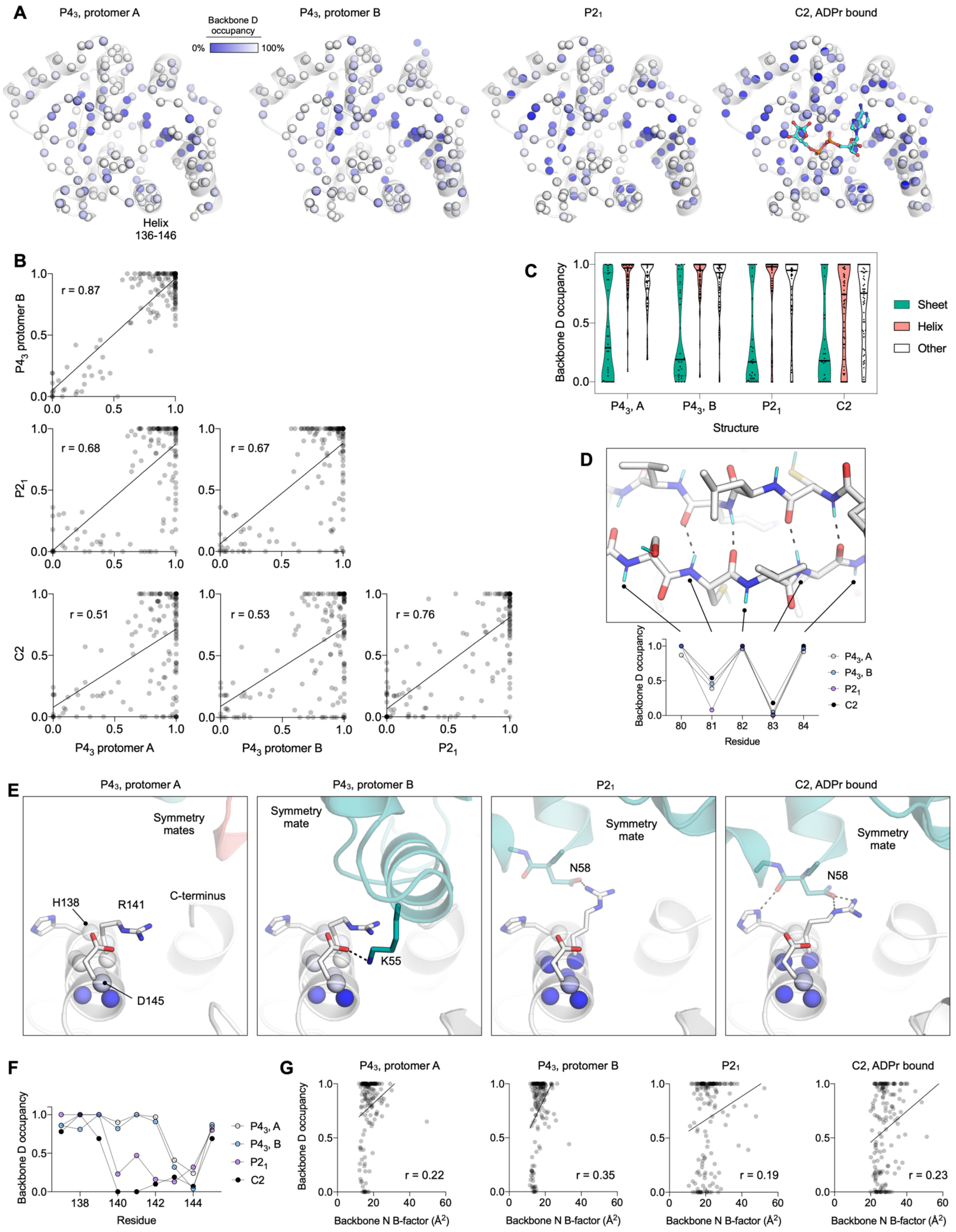
A
P4₃, protomer A
Backbone D occupancy
0%
100%
P4₃, protomer B
P2₁
C2, ADPr bound
Helix 136-146
B
r = 0.87
P4₃ protomer B
r = 0.68
r = 0.67
P2₁
r = 0.51
r = 0.53
r = 0.76
C2
P4₃ protomer A
P4₃ protomer B
P2₁
C
Backbone D occupancy
Sheet
Helix
Other
P4₃, A
P4₃, B
P2₁
C2
Structure
D
Backbone D occupancy
Residue
E
P4₃, protomer A
Symmetry mates
H138
R141
C-terminus
D145
P4₃, protomer B
Symmetry mate
K55
P2₁
Symmetry mate
N58
C2, ADPr bound
Symmetry mate
N58
F
Backbone D occupancy
Residue
G
P4₃, protomer A
r = 0.22
P4₃, protomer B
r = 0.35
P2₁
r = 0.19
C2, ADPr bound
r = 0.23
Backbone N B-factor (Å²)

**Fig. S4.** Comparison of backbone amide hydrogen-deuterium exchange across the three Mac1 neutron structures. (**A**) Cartoon representation of Mac1 showing the backbone nitrogens as spheres colored by backbone nitrogen deuterium occupancy (blue=0%, white=100%). (**B**) Plots showing the correlation between the refined occupancy of backbone deteriums in the three neutron structures. The gray line shows a linear fit of the data using GraphPad Prism. (**C**) Plot showing refined backbone D occupancy as a function of protein secondary structure. Secondary structure was calculated using the DSSP program (*98*). (**D**) Alternating high and low backbone D occupancy in a solvent exposed β-strand in the three Mac1 neutron structures. Occupancy varies from ~100% for the exposed backbone amides to 0-50% for the internally oriented amides. (**E**) Crystal contacts around the α-helix composed of residues 138-145 vary between the two protomers in the $P4_3$ structure, and the $P2_1$/C2 structures. Backbone nitrogens are shown with blue spheres, colored by D occupancy in the same way as (**A**). Hydrogen bonds between symmetry mates are shown as dashed black lines. (**F**) Plot showing the backbone D occupancy of the helix composed of residues 138-145. (**G**) Plot showing backbone D occupancy as a function of backbone N B-factor. The gray lines show a linear fit of the data using GraphPad Prism.

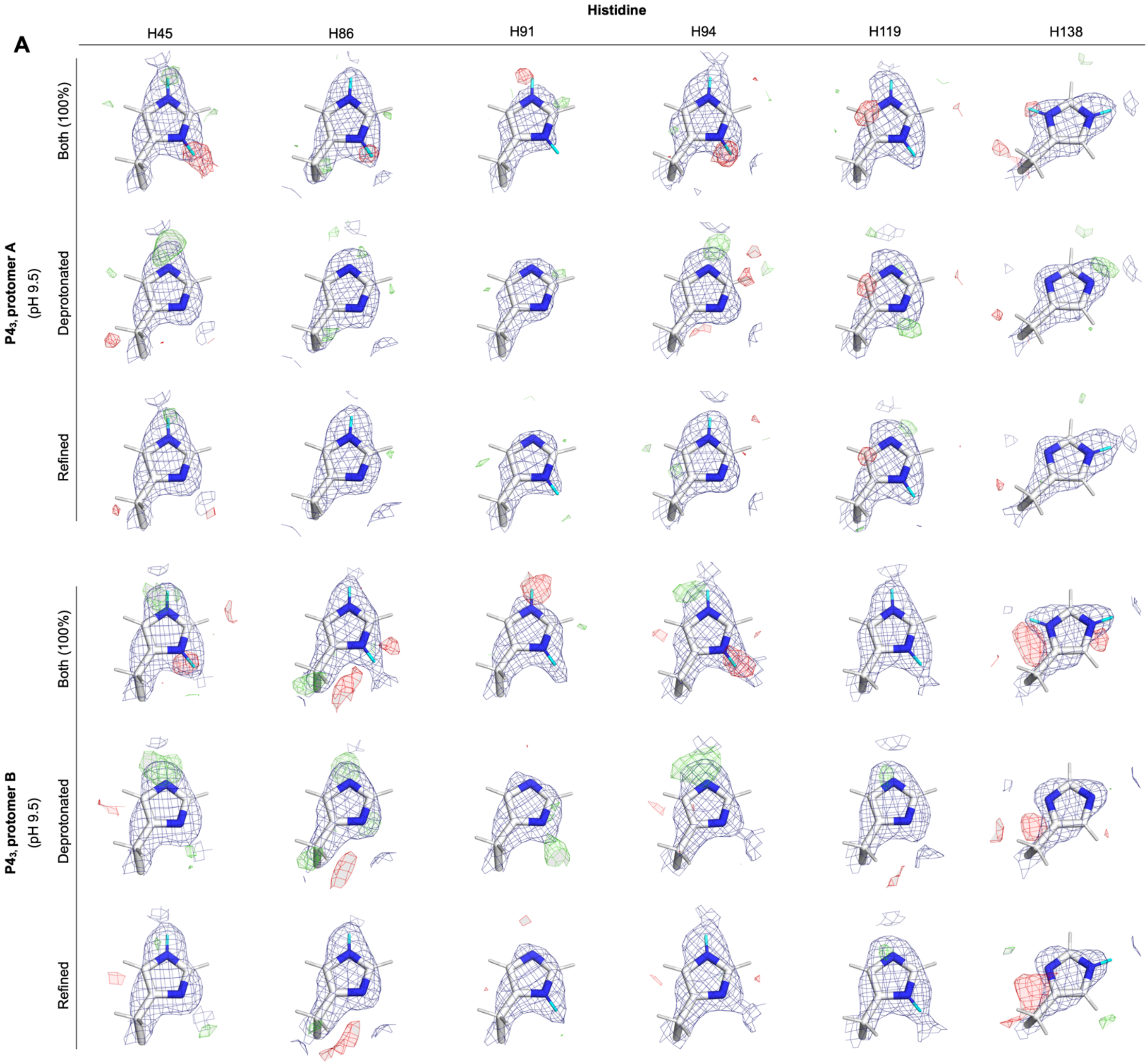
A
Histidine
H45
H86
H91
H94
H119
H138
P4₃, protomer A
(pH 9.5)
Both (100%)
Deprotonated
Refined
P4₃, protomer B
(pH 9.5)
Both (100%)
Deprotonated
Refined

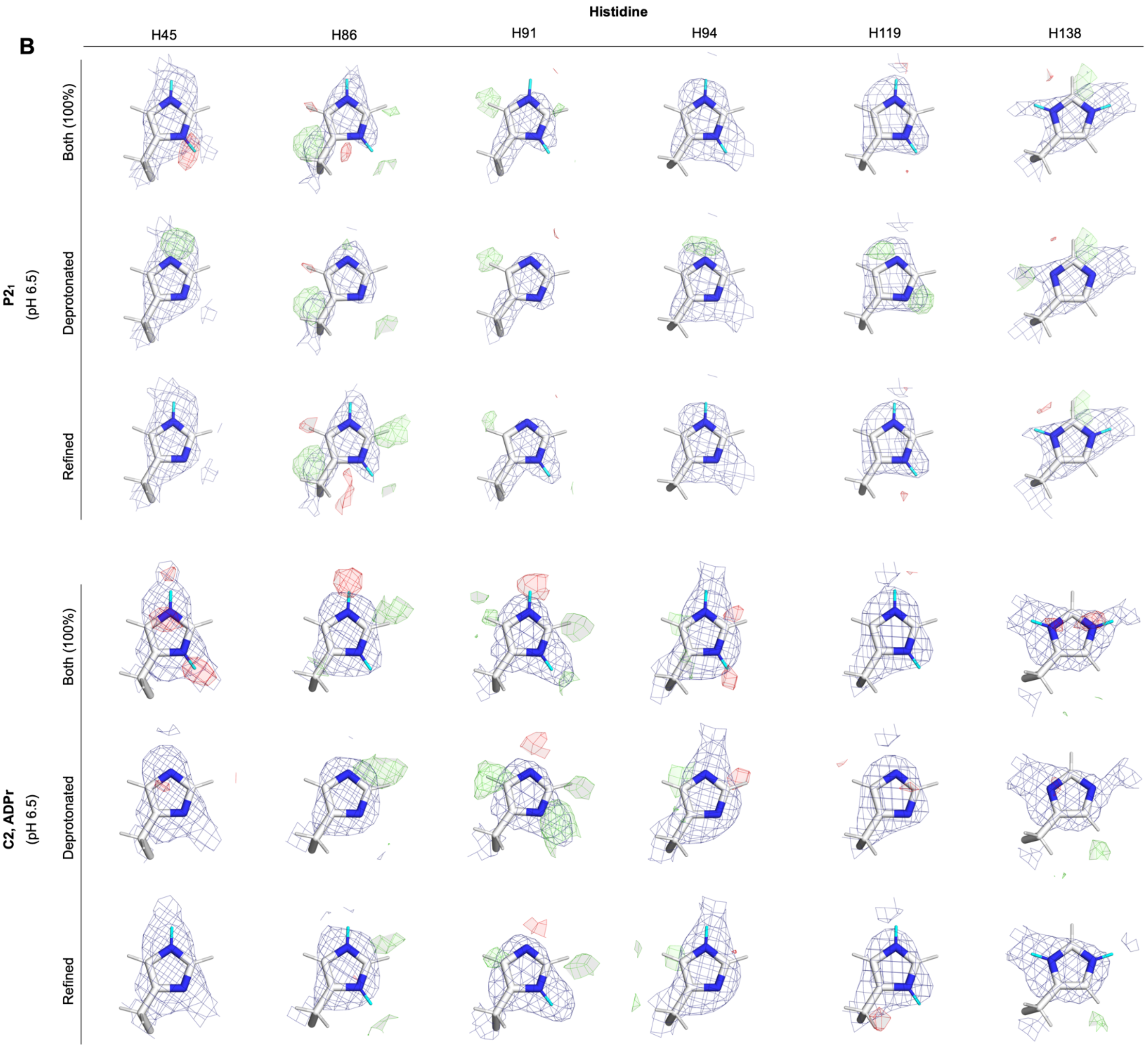

B
Histidine
H45
H86
H91
H94
H119
H138
$P2_1$
(pH 6.5)
C2, ADPr
(pH 6.5)
Both (100%)
Deprotonated
Refined
Both (100%)
Deprotonated
Refined

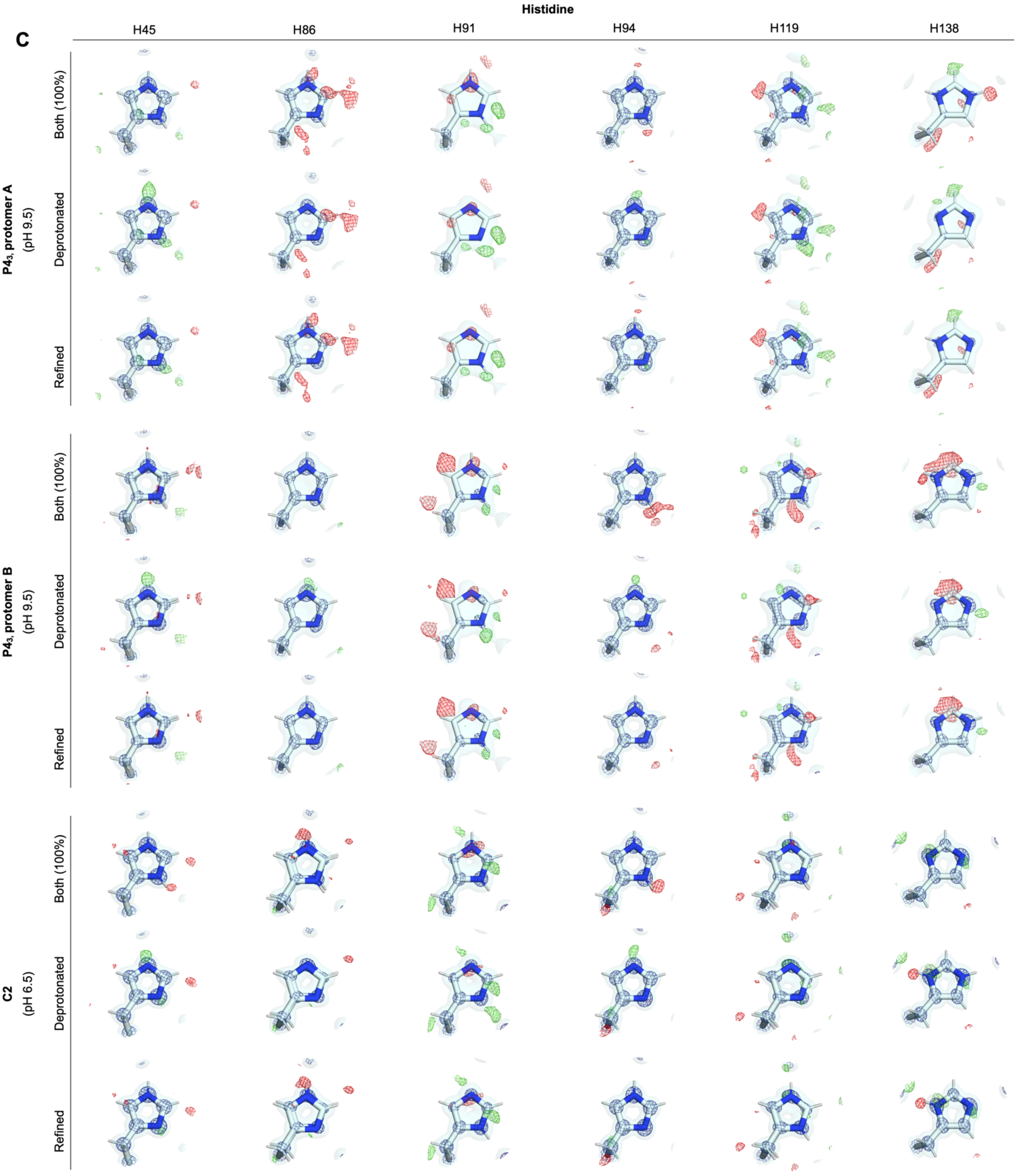

C
Histidine
H45
H86
H91
H94
H119
H138
P4₃, protomer A
(pH 9.5)
P4₃, protomer B
(pH 9.5)
C2
(pH 6.5)
Both (100%)
Deprotonated
Refined

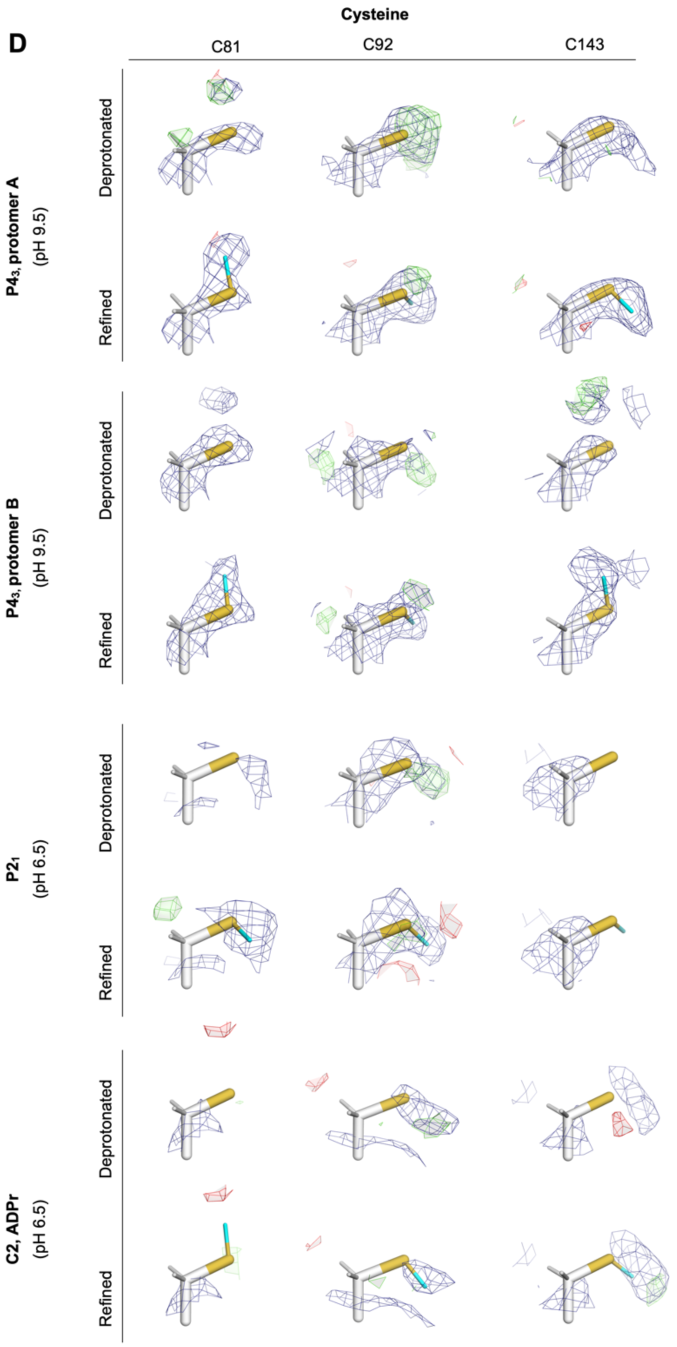


**Fig. S5.** Assignment of histidine and cysteine protonation states based on NSL / electron density maps. (**A**) NSL density maps used to assign the protonation states of histidine residues in the $P4_3$ crystal. The $2mF_O$-$DF_C$ (unfilled) NSL density map (blue mesh contoured at 1.5 σ) and the $mF_O$-$DF_C$ density map (green/red mesh contoured at ±2.5 σ) are shown calculated after: 1) refinement with Nδ1/Nε2 deuterium atoms set to 100% occupancy, 2) refinement prior to adding deuteriums to the Nδ1/Nε2 side chain atoms, and 3) refinement of the final model. (**B**) Same as (**A**) but showing maps calculated from the $P2_1$ and C2 structures. (**C**) Electron density maps showing evidence for histidine protons in the previously published 0.85 Å $P4_3$ structure (PDB code 7KQO) and the 0.77 Å C2 structure (PDB code 7KR0). The $2mF_O$-$DF_C$ electron density maps (blue mesh contoured at 4 σ, blue surface contoured at 1 σ) and the $mF_O$-$DF_C$ electron density maps (green/red mesh contoured at ±2.5 σ) are shown calculated after: 1) refinement of the doubly protonated form, 2) refinement of the doubly deprotonated form, and 3) refinement of the deposited coordinates. (**D**) NSL density maps for cysteine residues in Mac1. The $2mF_O$-$DF_C$ (unfilled) NSL density map (blue mesh contoured at 1.5 σ) and the $mF_O$-$DF_C$ density map (green/red mesh contoured at ±2.5 σ) are shown calculated after: 1) refinement prior to adding deuteriums to the Sγ side chain atom, and 2) refinement of the final model.

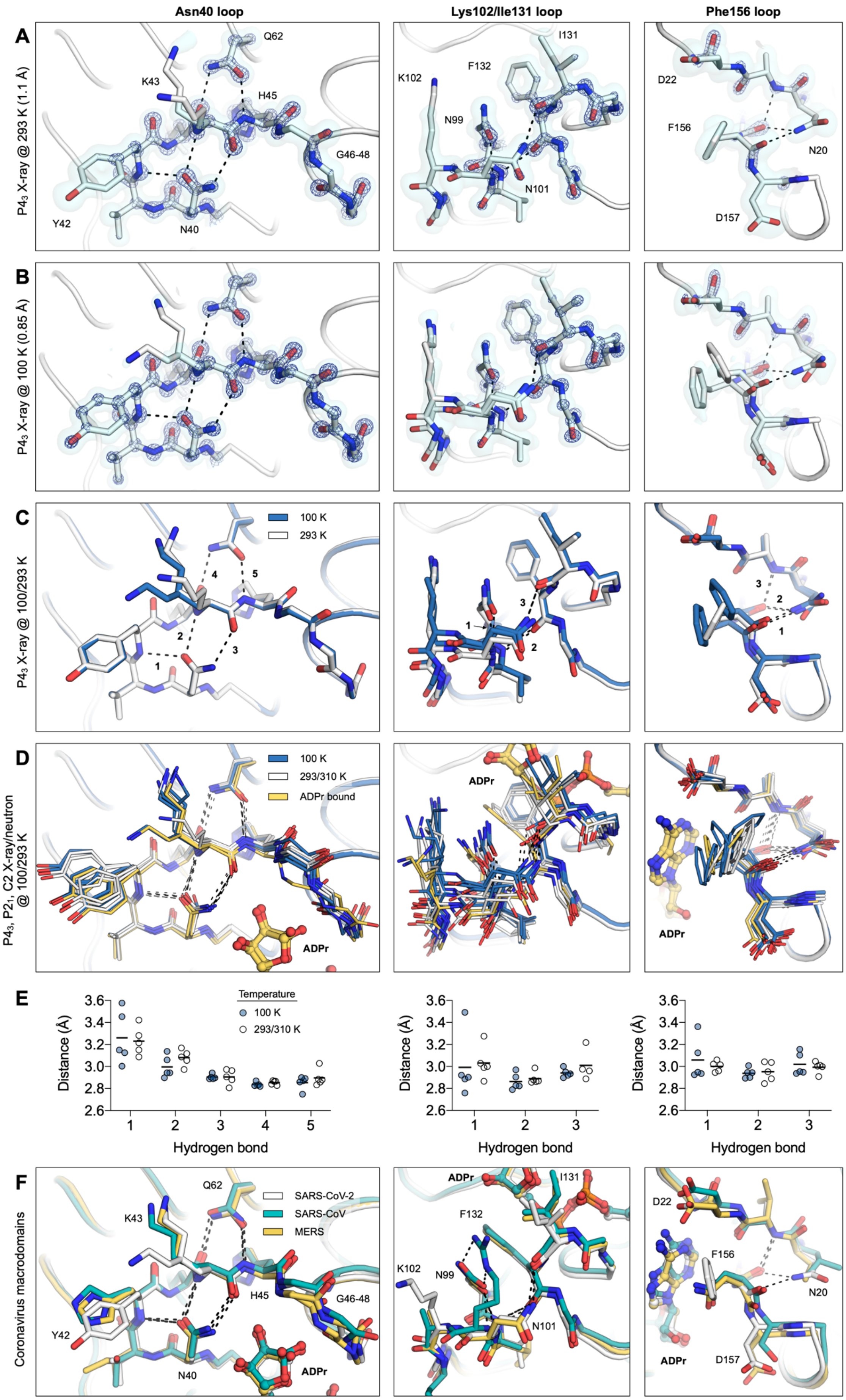
Asn40 loop
Lys102/Ile131 loop
Phe156 loop
A
P4₃ X-ray @ 293 K (1.1 Å)
Q62
K43
H45
G46-48
Y42
N40
I131
F132
K102
N99
N101
D22
F156
N20
D157
B
P4₃ X-ray @ 100 K (0.85 Å)
C
P4₃ X-ray @ 100/293 K
100 K
293 K
D
P4₃, P2₁, C2 X-ray/neutron @ 100/293 K
293/310 K
ADPr bound
ADPr
E
Distance (Å)
Temperature
100 K
293/310 K
Hydrogen bond
F
Coronavirus macrodomains
SARS-CoV-2
SARS-CoV
MERS

**Fig. S6.** Flexibility and hydrogen bond networks in the Mac1 active site. (**A**) The three panels show the Asn40 loop, the Lys102/Ile131 loop and the Phe156 loop for the 1.1 Å $P4_3$ X-ray structure determined at 293 K (protomer A, PDB code 7TWH). The $2mF_O$-$DF_C$ electron density map is shown (blue mesh/surface contoured at 4/1 σ). Hydrogen bonds are shown with dashed black lines. The protein is shown with white sticks/surface. For clarity, only selected side chains are shown. (**B**) Same as (**A**) but showing the 0.85 Å $P4_3$ X-ray structure determined at 100 K (protomer A, PDB code 7KQO). (**C**) Alignment of structures shown in (**A**) and (**B**). Hydrogen bonds are numbered according to the plot of distances shown in (**E**). (**D**) Alignment of the three X-ray/neutron structures reported in this work (PDB code 7TX3, 7TX4, 7TX5) with the 1.1 Å X-ray structure at determined at 293 K (P43, PDB code 7TWH) and the previously reported apo and ADPr-bound $P4_3$ structures (PDB code 7KQO/7QKP) and the C2 structures at 100 and 310 K (PDB codes 7KR0 and 7KR1). (**E**) Plot showing hydrogen bond lengths from the structural ensemble shown in (**E**). Hydrogen bond numbering is shown in (**D**). (**F**) Alignment of the NSP3 macrodomain from SARS-CoV-2 (PDB code 7QKP) with the SARS-CoV (PDB code 2FAV) and MERS (PDB code 5HOL) macrodomains.

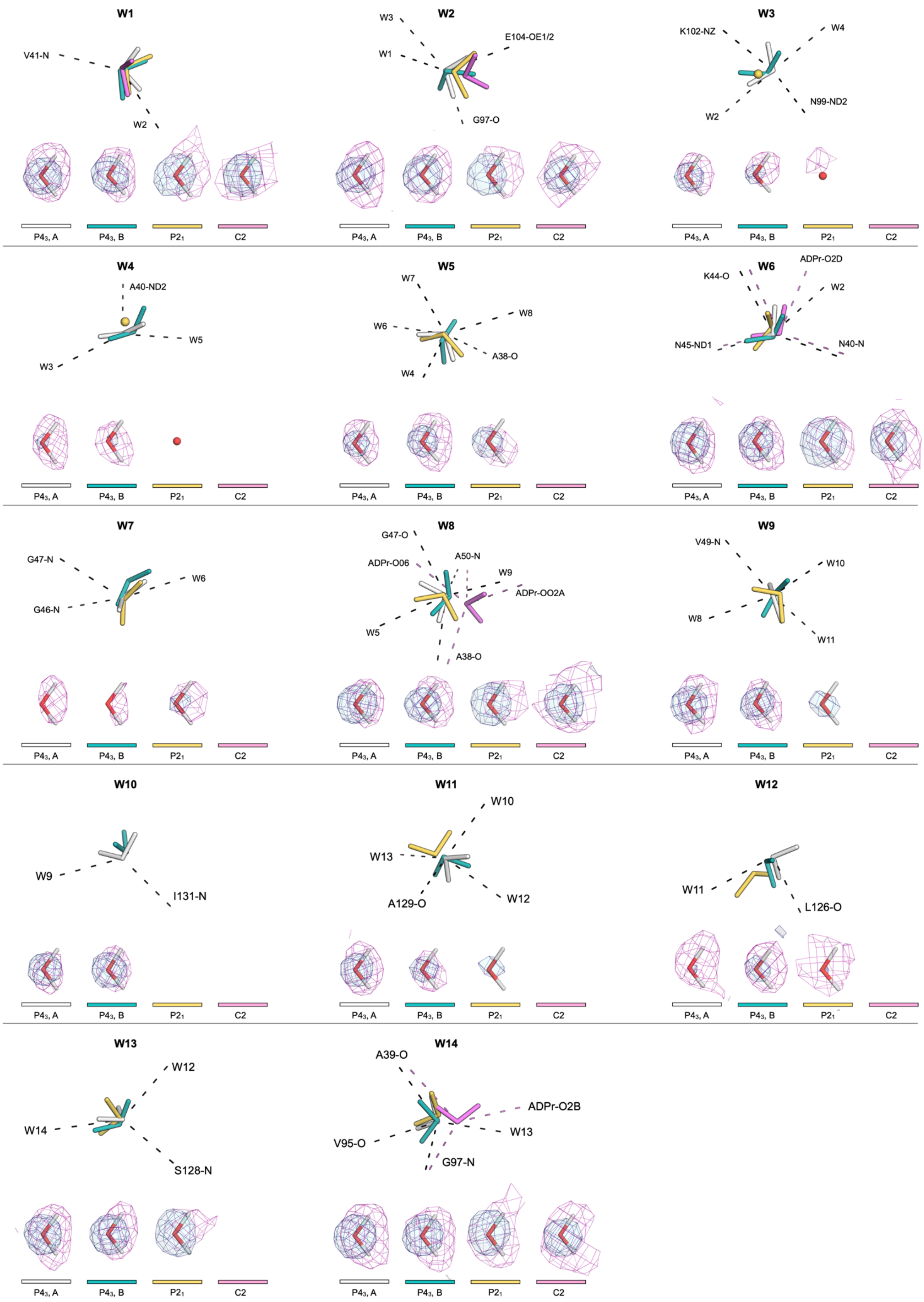
W1
V41-N
W2
W2
W3
W1
E104-OE1/2
G97-O
W3
K102-NZ
W4
N99-ND2
W2
W4
A40-ND2
W5
W3
W5
W7
W8
W6
A38-O
W4
W6
ADPr-O2D
K44-O
W2
N45-ND1
N40-N
W7
G47-N
W6
G46-N
W8
G47-O
A50-N
ADPr-O06
W9
ADPr-OO2A
W5
A38-O
W9
V49-N
W10
W8
W11
W10
W9
I131-N
W11
W10
W13
A129-O
W12
W12
W11
L126-O
W13
W12
W14
S128-N
W14
A39-O
ADPr-O2B
V95-O
W13
G97-N
P4₃, A
P4₃, B
P2₁
C2

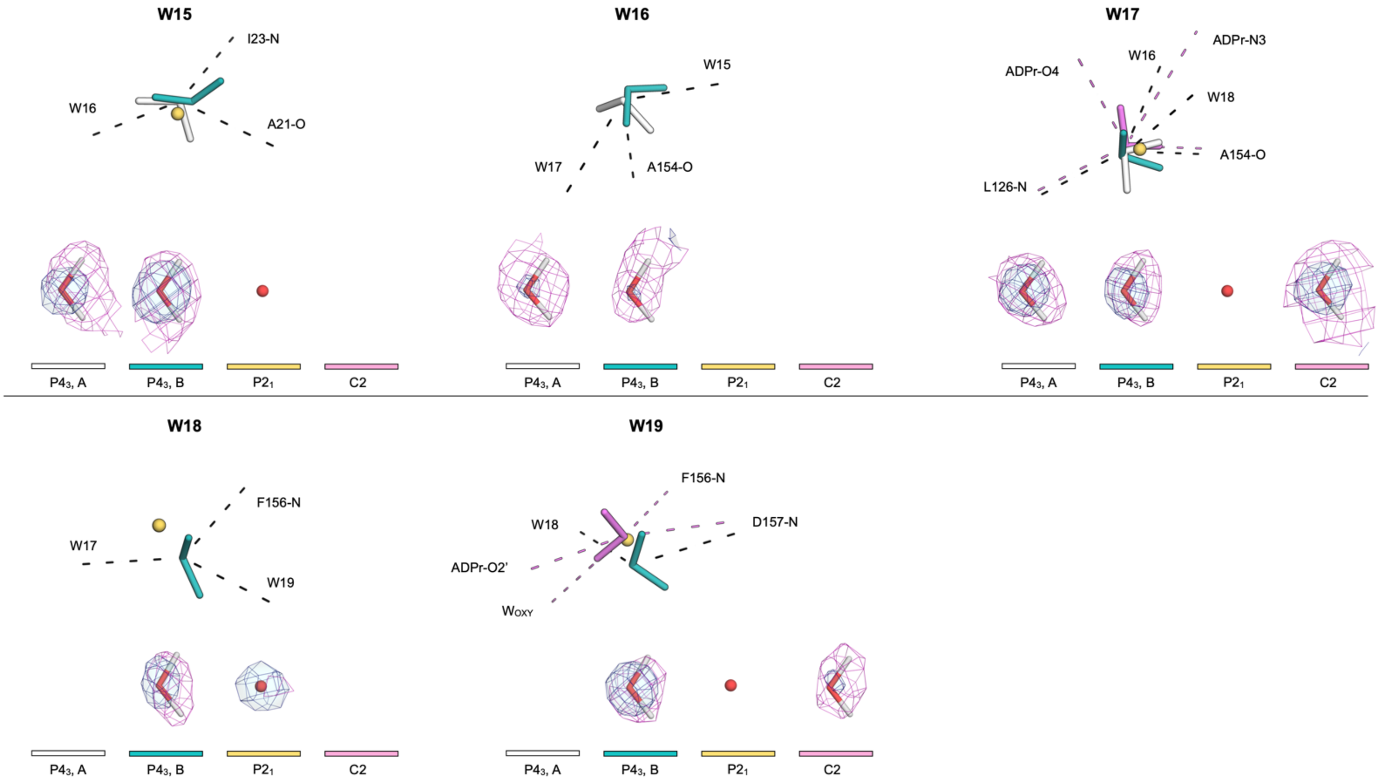


**Fig. S7.** Evidence for $D_2O$ orientations from neutron scattering length density maps. The 2m$F_O$-D$F_C$ (unfilled) NSL density maps (purple mesh contoured at 2 σ) and the 2m$F_O$-D$F_C$ electron density maps (blue mesh/surface contoured at 2 σ) are shown for the $P4_3$, $P2_1$ and C2 neutron structures (PDB codes 7TX3, 7TX4, 7TX5). Hydrogen bonds to neighboring residues are shown with dashed black lines. For the C2 structure, ADPr specific hydrogen bonds are shown with pink dashed lines. For $D_2O$ molecules that were modeled without deuterium atoms, the oxygen is shown as a red sphere.

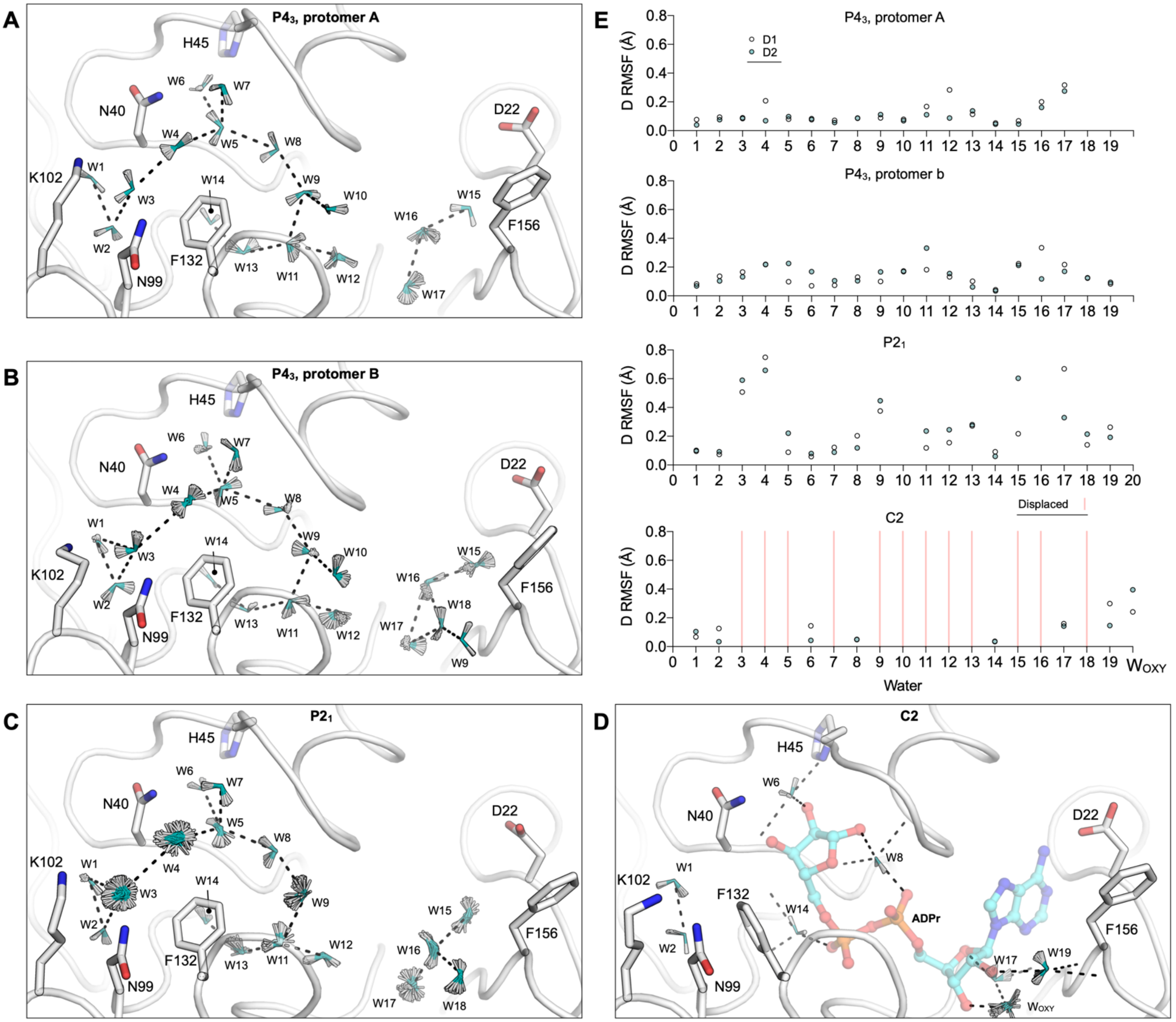


**Fig. S8.** Assessing the variability in water orientations in joint X-ray/neutron refinement with phenix.refine. (**A**-**D**) Alignment of the 100 structures after refinement starting from coordinates where the $D_2O$ deuterium atoms were randomly shifted by 0.5 Å. $D_2O$ molecules are shown with sticks with the oxygen colored teal and the deuterium atoms colored white. For clarity, only selected side chains are shown. (**E**) Plots showing RMSF values for $D_2O$ deuteriums calculated across the structures shown in (**A**-**D**).

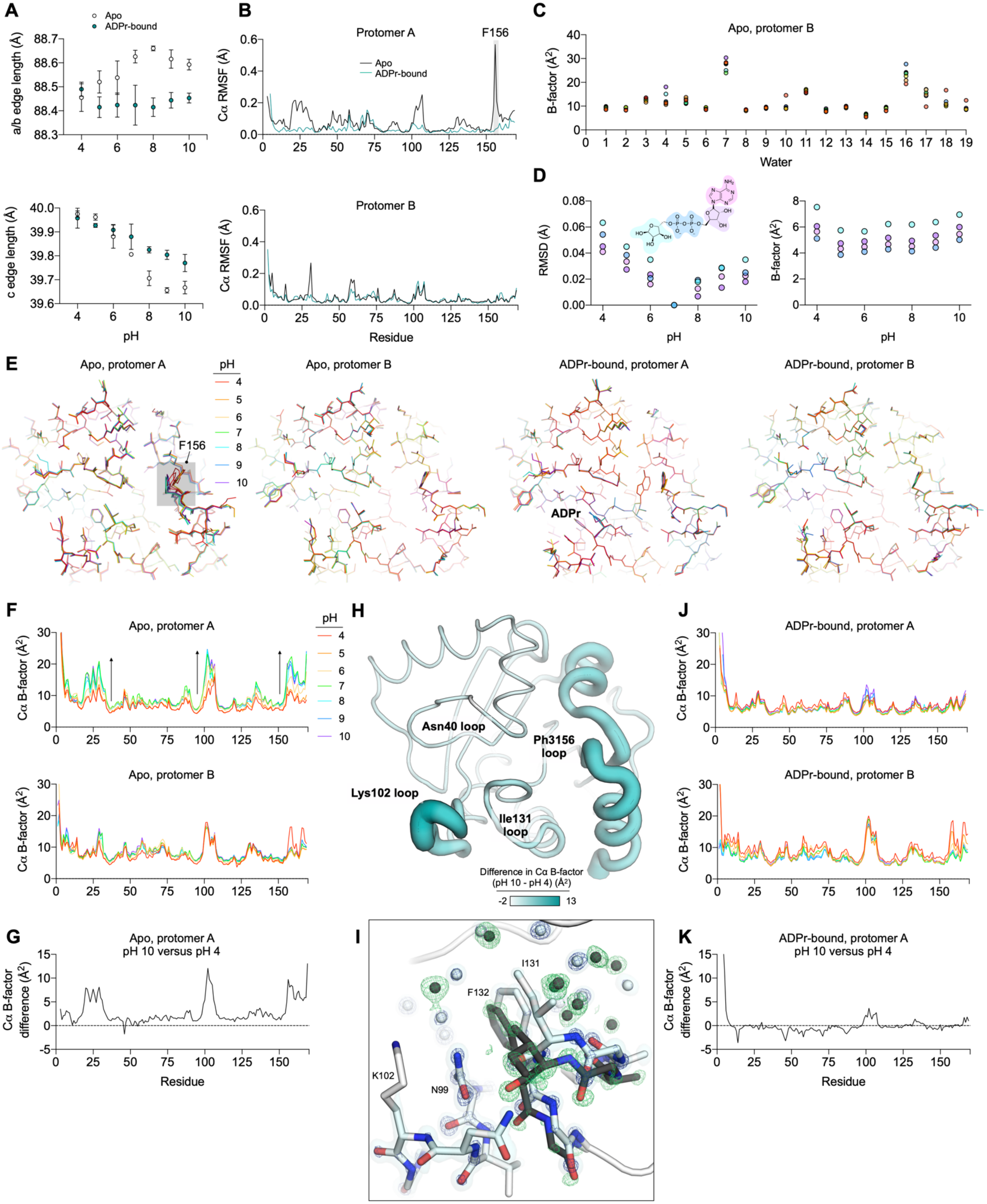
A
Apo
ADPr-bound
a/b edge length (Å)
c edge length (Å)
pH
B
Protomer A
F156
Apo
ADPr-bound
Cα RMSF (Å)
Protomer B
Residue
C
Apo, protomer B
B-factor (Å²)
Water
D
RMSD (Å)
B-factor (Å²)
pH
E
Apo, protomer A
Apo, protomer B
ADPr-bound, protomer A
ADPr-bound, protomer B
F156
ADPr
F
Apo, protomer A
Apo, protomer B
Cα B-factor (Å²)
G
Apo, protomer A
pH 10 versus pH 4
Cα B-factor difference (Å²)
Residue
H
Asn40 loop
Ph3156 loop
Lys102 loop
Ile131 loop
Difference in Cα B-factor (pH 10 - pH 4) (Å²)
I
I131
F132
K102
N99
J
ADPr-bound, protomer A
ADPr-bound, protomer B
Cα B-factor (Å²)
K
ADPr-bound, protomer A
pH 10 versus pH 4
Cα B-factor difference (Å²)
Residue

**Fig. S9.** Investigating Mac1 conformational diversity with pH-shift crystallography. (**A**) Plots showing the pH-dependent change in the $P4_3$ unit cell. Each data point represents the unit cell measurements from at least three crystals, with error bars showing ± the standard deviation. (**B**) Plots showing Cα RMSF calculated across the seven apo Mac1 structures from pH 4 to 10. (**C**) Plot showing the B-factors for water molecules in the active site of protomer B for the seven apo Mac1 structures from pH 4 to 10. (**D**) Right: plot showing RMSD values for the different moieties of ADPr as a function of pH. RMSD values were calculated relative to the structure at pH 7. Left: ADPr B-factors as a function of pH. (**E**) Alignment of the seven Mac1 structures from pH 4 to 10. Structures are colored by pH. (**F**) Plot showing Cα B-factors of the apo Mac1 structures from pH 4 to 10. (**G**) Plot showing the difference in Cα B-factor between pH 10 and 4 for the apo Mac1 structures. (**H**) Difference in B-factors shown in (**G**) mapped onto the Mac1 structure. (**I**) Evidence for an alternative conformation of the Ile131 loop in the structure determined from Mac1 crystals soaked at pH 4. The major conformation is shown with white sticks/spheres/cartoon with the corresponding $2mF_O\text{-}DF_C$ electron density map (blue mesh/surface contoured at 4/1 σ). The alternative conformation is shown with black sticks/spheres with the corresponding $mF_O\text{-}DF_C$ electron density map (green mesh contoured at +3 σ). (**J**) Plot showing Cα B-factors of the ADPr-bound Mac1 structures determined from pH 4 to 10. Coloring is the same as (**F**). (**K**) Plot showing the difference in Cα B-factor between pH 10 and 4 for the ADPr-bound Mac1 structures.

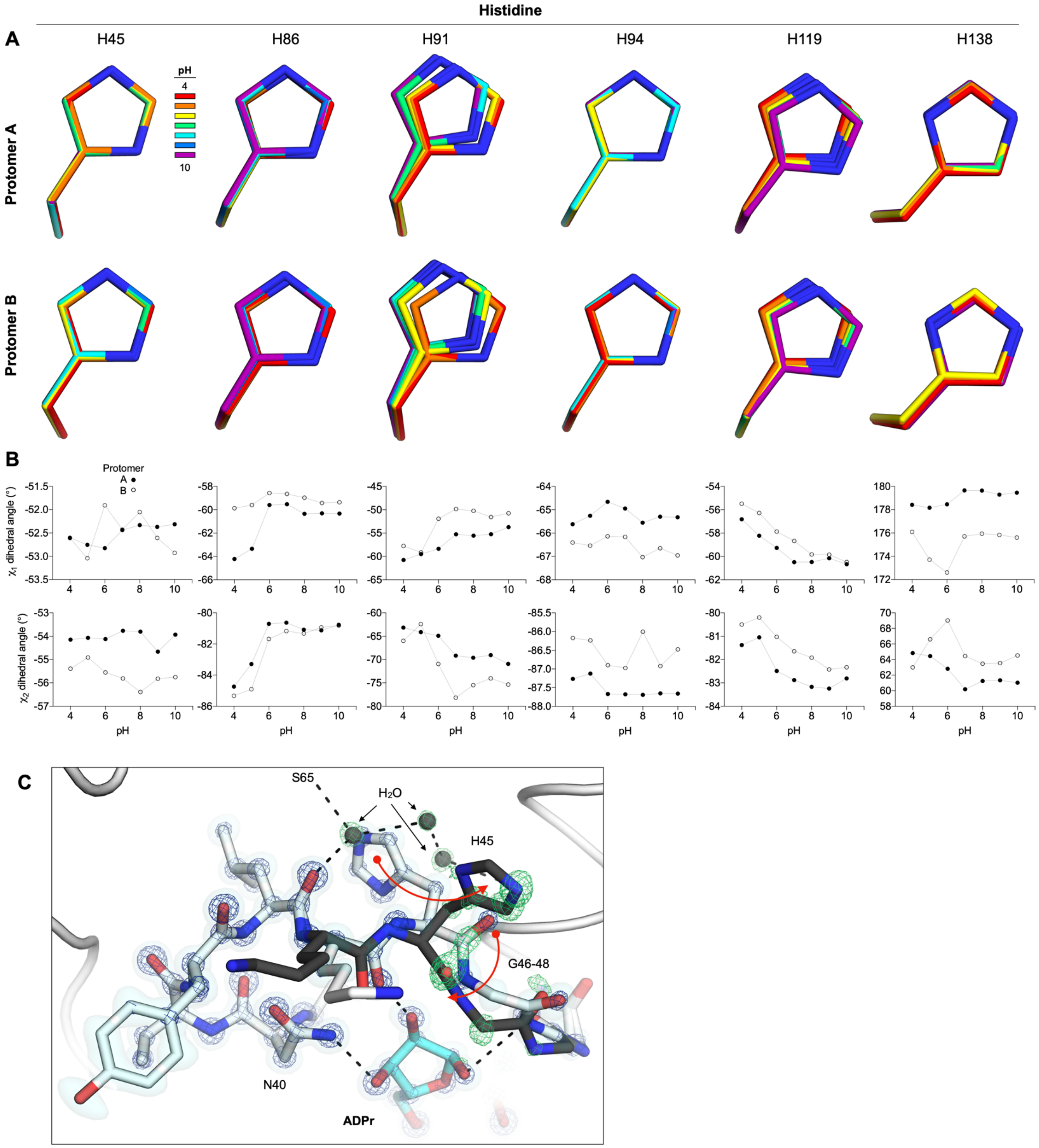


**Fig. S10.** pH-induced conformational changes in Mac1 histidines. (**A**) Alignment of Mac1 histidines from the apo structures determined from pH 4 to 10. (**B**) Plot showing the $\chi_1$ and $\chi_2$-dihedral angles for the histidines shown in (**A**). (**C**) Evidence for an alternative conformation of His45 and the Gly46-48 loop in the Mac1 structure with ADPr-bound determined at pH 4. Red arrows indicate the conformational changes. The major conformation is shown with white sticks/spheres/cartoon with the corresponding $2mF_O$-$DF_C$ electron density map (blue mesh/surface contoured at 4/1 σ). The alternative conformation is shown with black sticks/spheres with the corresponding $mF_O$-$DF_C$ electron density map (green mesh contoured at +3 σ).
